# Tumor PD-L1 selectively suppresses type I interferon in myeloid cells to suppress CTL recruitment to promote lung metastasis

**DOI:** 10.1101/2021.06.18.449028

**Authors:** John D. Klement, Priscilla S. Redd, Chunwan Lu, Alyssa D. Merting, Dakota B. Poschel, Dafeng Yang, Gang Zhou, David H. Munn, Kebin Liu

## Abstract

The mechanism underlying tumor cell PD-L1 (tPD-L1) induction of immune suppression through T cell PD-1 is well-known, but the mechanism underlying tPD-L1 induction of immune suppression via an intermediate cell is incompletely understood. We report here that tPD-L1 does not suppress cytotoxic T lymphocyte (CTL) activation and lytic function when only tumor cells and CTLs are present. Strikingly, knocking out PD-L1 in tumor cells has no effect on primary tumor growth, but significantly decreases lung metastasis in a CTL-dependent manner. Depletion of myeloid cells impaired tPD-L1 promotion of lung metastasis. Single-cell RNA sequencing revealed that tPD-L1 engages myeloid PD-1 (mPD-1) to antagonize type I interferon (IFN-I) and STAT1 signaling to repress *Cxcl9* and *Cxcl10* expression to impair CTL recruitment to lung metastases. Human patient response to PD-1 blockade immunotherapy correlates with IFN-I response in myeloid cells. Our data determines that the tPD-L1/mPD-1/IFN-I/STAT1/Cxcl9/10 axis controls CTL tumor infiltration in lung metastasis.

## Introduction

PD-L1 and PD-1 immune checkpoint blockade (ICB) immunotherapy is believed to function by blocking the engagement of PD-L1 to co-inhibitory receptor PD-1 expressed by activated and exhausted cytotoxic T lymphocytes (CTLs) ^1^. PD-1 engagement on CTLs has been shown to result in decreased activation and effector functions due to SHP-2 mediated dephosphorylation of the T cell receptor and CD28 signaling cascades, driving CTLs toward an exhausted fate ^2^. PD-1/L1 blockade re activates and expands the tumor-reactive CTLs, leading to tumor suppression ^3^. Particular attention has been paid to the role of tumor-expressed PD-L1 (tPD-L1), which has been hypothesized to serve as a direct molecular shield against CTL lysis of tumor cells ^4^.

Emerging experimental data has started to reveal alternative pathways to this paradigm, and suggest that response of CTLs to PD-1/L1 blockade may depend on tumor anatomic sites and occur principally in other sites, rather than the primary tumor site ^5^. A defining hallmark of PD-1/L1 ICB immunotherapy therapy is the sustained remission of disease experienced by responders ^6^. As the dominant cause of mortality is metastatic growth ^7^, the durable survival benefit conferred by PD-1/L1 blockade thus arises from coordinated suppression of metastases ^8^. Recognition of this fact has led to increased interest in PD-1/L1 ICB in the neoadjuvant and the adjuvant setting ^9, 10^. Neoadjuvant and adjuvant therapy is administered at an earlier point in disease course, either before or immediately following primary tumor resection, respectively. In doing so, these therapies extend survival by containing metastases ^8, 11^. Robust tumor expression of PD-L1 has been shown to promote both micro- and macro-metastatic tumor growth, with enhancement at metastatic tumor sites as compared to primary tumor sites ^12, 13^. However, how tPD-L1 might differentially drive metastasis from primary tumor is unknown.

Emerging experimental data over the past decade has highlighted the importance of PD-1 expression in other cell populations, including myeloid, dendritic, and tumor cells, in tumor immune evasion ^14, 15, 16^. However, how non-T cell-expressed PD-1 connects tPD-L1 to CTL suppression is incompletely understood. We report here that tPD-L1 exhibits no direct effect on CTL activity when only tumor cells and tumor-specific CTLs are present. We show instead that tPD-L1 targets myeloid cell-expressed PD-1 (mPD-1) to suppress type I IFN expression and function. This represses Cxcl9 and Cxcl10 expression in myeloid cells and thereby impairs CTL recruitment to lung niche to enhance established lung metastasis in a manner independent of promoting primary tumor growth. Our data reveals a critical role for mPD-1 in bridging tPD-L1 to CTL recruitment in tumor metastasis.

## Results

### Tumor PD-L1 exhibits no direct protection of tumor cells from CTL cytotoxicity

To determine whether tPD-L1 protects tumor cells from CTL killing when only target tumor cells and tumor-specific CTLs are present in the microenvironment, we deleted *Cd274*, which encodes PD-L1, in 4T1 (Fig S1A-C), a murine triple negative breast cancer (TNBC) cell line, and CT26, a murine mismatch repair proficient (MMRp) colorectal cancer (CRC) cell line. The choice was made to mimic TNBC and MMRp CRC, which show minimal response to ICB immunotherapy in clinically advanced disease ^17, 18^. Loss of PD-L1 renders cell lines unable to engage CTL-expressed PD-1, as they show no expression of PD-L2, the only alternative ligand for PD-1 (Fig S1D).

To directly investigate the functional consequences of PD-1:tPD-L1 interaction, we utilized an in vitro CTL-tumor interaction system. 2/20 is a tumor-specific CTL line sensitive to PD-1 mediated inhibition (Fig S1E) that recognizes the H2-L^d^-restricted AH1 epitope (residues 6-14) of gp70 protein endogenously expressed by 4T1 and CT26 cells ^19^. As with effector CTLs, 2/20 cells exert antigen-specific cytotoxicity within minutes and do not need to undergo differentiation from a naïve state before acquiring effector function (unlike naïve transgenic T cells, i.e. OT-1, Pmel-1), allowing for recapitulation of the effector CTL:tumor interaction with high fidelity. This tumor model mimics the tPD-L1:CTL PD-1 interaction in the effector phase in the tumor microenvironment. Furthermore, 2/20 are directed against an endogenously expressed antigen – rather than artificially over-expressed at supra-physiological levels – that undergoes silencing and thus effectively models a tumor-expressed antigen (Fig 1A)^20^.

**Figure 1.**
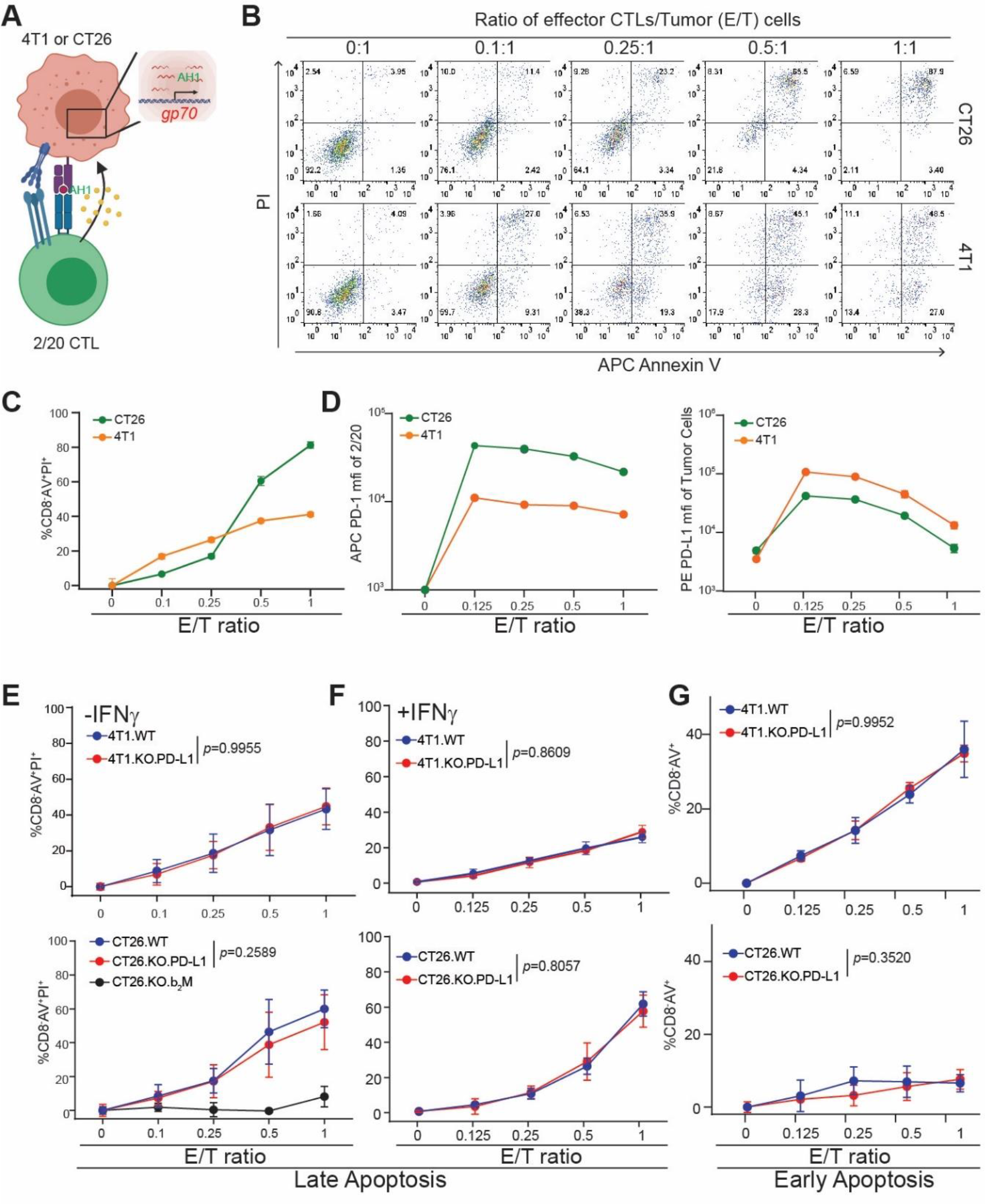
tPD-L1 does not confer direct protection from CTL killing. **A)** Schematic of co-culture system. **B)** Representative images of tumor cell apoptosis following co-culture with 2/20 CTLs at indicated ratios. Plots are gated on CD8α^−^ cells. Live cells: AnnexinV^−^PI^−^; Early apoptotic cells: AnnexinV^+^PI^−^; Late apoptotic cells: AnnexinV^+^PI^+^. **C)** Quantification of cell death of target cells (CD8^−^AV^+^PI^+^) by flow cytometry following co-culture. N = 3/condition. Mean ± SE **D)** Surface protein expression following overnight co-culture at indicated Effector/Target (E/T) ratios. N=3/condition. Mean ± SE. **E-F)** Cell death after overnight co-culture following no pretreatment (E) or pretreatment with IFNγ (F). N=6/condition. **G)** Proportion of early apoptotic cells after four hours of co-culture at indicated ratios. N=4/condition. Two-way ANOVA with Sidak’s multiple comparison.

2/20 CTLs displayed dose-dependent cytotoxicity against target cell lines (Fig 1B-C, S1H). Co-culture enhanced PD-1 expression level in 2/20 CTLs and PD-L1 expression on tumor cells. The degree of PD-L1/PD-1 expression varied with initial E/T input, allowing us to assay differing ratios of PD-1 to PD-L1 expression (Fig 1D). Surprisingly, and contradicting the hypothesis of tPD-L1 as a molecular shield, tPD-L1 expression conferred no protection against 2/20 cytotoxicity in the 4T1 or CT26 at early or late time points. This phenomenon was not rescuable by pretreatment with IFNγ to elevate surface PD-L1 expression (Fig 1E-G, S1G-H). Knocking out β-2-microglobulin (B2M) in tumor cells abolished CTL function in killing the tumor cells, validating an antigen-specific killing of tumor cells by the 2/20 CTLs (Fig. 2E). These results indicate that tPD-L1 confers no survival advantage when an antigen-specific effector CTL encounters a tumor cell bearing a cognate peptide-MHC complex, free from any other tumor microenvironmental factors.

**Figure 2.**
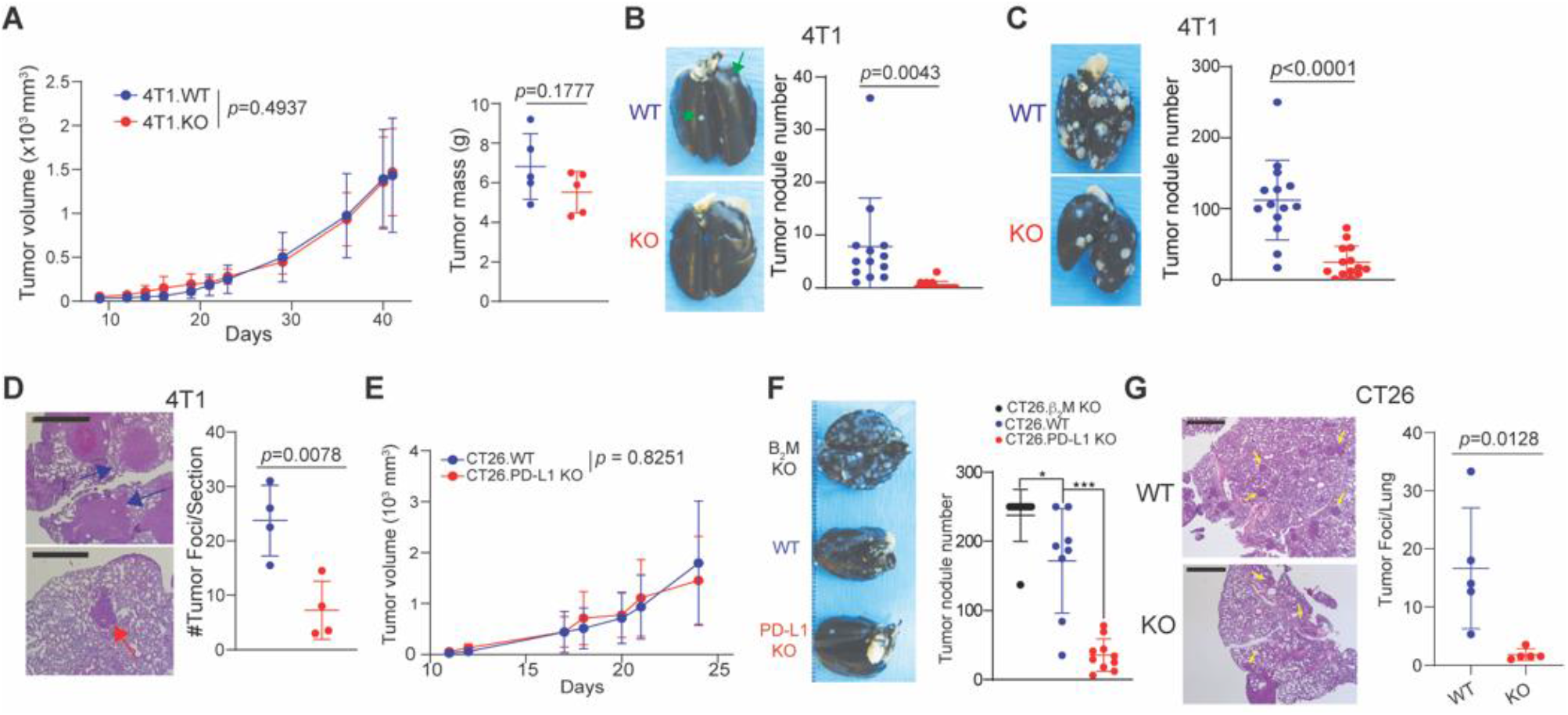
tPD-L1 selectively enhances metastasis independently of primary tumor growth. **A)** Growth curves (left) and final mass (right) of 4T1 WT and PDL1-KO following orthotopic injection. Representative of three independent experiments. N= 5-15/condition. 2-way ANOVA. 4T1 WT and PDL1-KO spontaneous metastasis to the lung following orthotopic injection. N= 5-10/condition, two independent experiments. 2-way ANOVA. **C-D)** Quantification of 4T1 experimental metastatic foci by visual (C, N=15/condition) and microscopic (D, N=4/condition) inspection. Scale: 1 mm **E)** Tumor growth curve of CT26 following subcutaneous injection. N=5/condition. **F-G)** Quantification of CT26 experimental metastatic foci by visual (F, N=10/condition) and microscopic (G, N=5/condition) inspection. Scale: 1 mm. All error bars in figure represent mean ± SD.

### Tumor PD-L1 enhances metastasis independent of primary tumor growth

We next sought to extend these in vitro findings to the in vivo setting. We first examined tumor growth in the site of tumor cell injection (primary tumor growth). Initial orthotopic injection of PD-L1 WT and PD-L1 KO 4T1 cells (Fig. S2C) to syngeneic mice led to tumor growth at the site of injection. Intriguingly, no difference was observed in primary tumor growth, supporting our in vitro cytotoxicity results (Fig 2A, S2B). No differences in tumor growth were observed in the WT and PD-L1 KO CT26 tumor at the site of tumor cell injection (Fig. 2E).

4T1 tumor cells give rise to spontaneous lung metastasis if orthotopically transplanted to the mammary gland. Orthotopic injection of a minimal dose of cells allowed for dissemination and growth of metastatic tumor in the lung without resection of the primary tumor (Fig S2A). We then examined spontaneous lung metastasis. Although loss of tPD-L1 does not affect primary tumor growth (Fig 2A & E), loss of tPD-L1 impaired spontaneous metastasis to the lung (Fig 2B). To extend this finding, we utilized an experimental lung metastasis model in which lung metastasis is quantified following tail vein injection (Fig S2D). Lung metastasis was observed in PD-L1-deficient 4T1 at both the gross and histological level (Fig 2C-D). To confirm these results were not 4T1- or TNBC-specific, we repeated these experiments in the MMRp CRC model CT26. Similarly, tPD-L1 promoted metastasis independent of primary tumor growth (Fig 2F-G).

The above observations suggest that tPD-L1 functioned either by promoting colonization of disseminated tumor cells (DTC) or by promoting growth of metastases in the lung.

Increased infiltration of CD3^+^ cells after loss of tPD-L1 in lung, but not in the primary tumor were observed (Fig. 3A), which indicate that tPD-L1 blocked T cell infiltration at the metastatic site. No difference in initial tumor cell colonization as measured by tumor cell gp70 level in the lungs was seen one day after tail vein injection (Fig. 3B), demonstrating that PD-L1 does not enhance DTC colonization efficiency in the lung. Similarly, PD-L1-deficient and competent 4T1 metastasized equally in adaptive-immune deficient congenic RAGII-KO mice (Fig 3C), indicating that tPD-L1 requires a functioning adaptive immune system to promote lung metastasis. Genetic ablation of MHC class I antigen presentation, as well as antibody-mediated depletion of CD8^+^ T cells, revealed that tPD-L1 drove metastasis by suppressing killing by CTLs (Fig 2F, 3D). Taken together, these results demonstrate that lung-resident DTCs and metastasis utilize tPD-L1 to evade CTL-mediated clearance, rather than driving extravasation or seeding from the primary tumor site.

**Figure 3.**
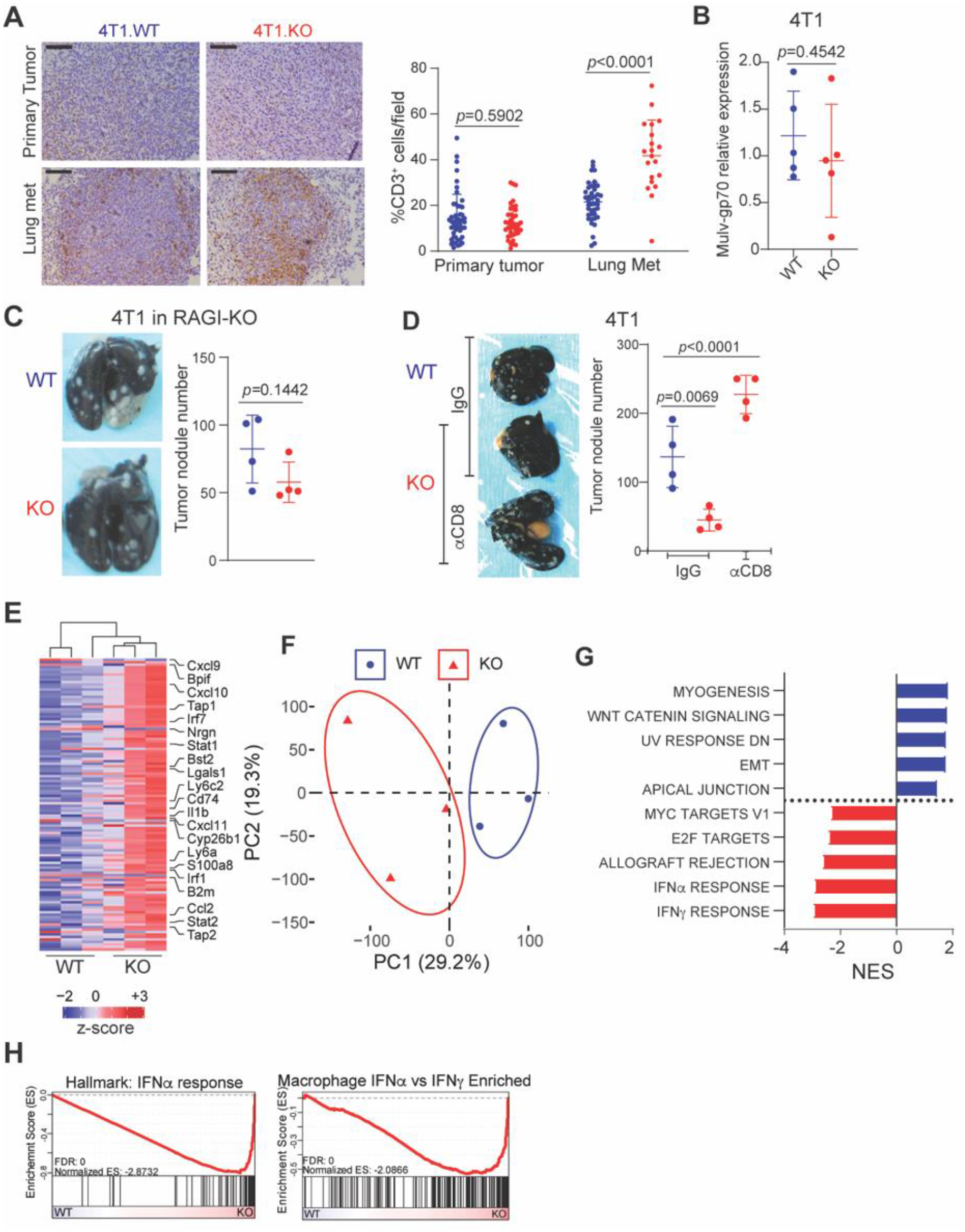
Loss of tPD-L1 limits metastasis by amplifying inflammatory and CTL-driven responses. **A)** Representative images (left) and quantification (right) of CD3 expression. Pooled from two independent experiments. N=8/condition. Scale: 1mm **B)** *Gp70* expression in lung twenty-four hours post-injection of indicated cell lines. Two independent experiments. N=5/condition. **C)** Representative image (left) and quantification (right) of experimental metastasis in Balb/c RAGII-KO N=4/condition, two independent experiments. **D)** Representative image (left) and quantification (right) of tumor nodules following CD8 depletion. N=4/condition. **E)** Heatmap of top 125 most variably expressed genes detected in lungs colonized by 4T1 WT or PDL1-KO. **F)** Principal component analysis. G) Top 5 enriched gene signatures from MSigDB Hallmarks signature set in WT (blue) or PDL1-KO (red) colonized lungs based on normalized enrichment score (NES). **H)** Enrichment plot of indicated signatures. All error bars in figure represent mean ± SD.

Given our observation showing that tPD-L1 does not directly inhibit CTL killing of 4T1 and CT26 (Fig. 1), we therefore hypothesized that a third factor, only present at lung metastatic niche, was necessary for tPD-L1 to exert its indirect protective role. To test this hypothesis and identify this third factor, we performed RNA sequencing of lungs colonized by WT and PD-L1-deficient 4T1. Differentially expressed transcripts include those involved in antigen presentation (*B2m, Cd74, Tap1)*, interferon response (*Stat1, Bst2, Irf1*) and CTL recruitment (*Cxcl9, Cxcl10 and Cxcl11*) (Fig 3E, S3A). Dimensionality reduction by principal component analysis separated on the first principal component (Fig 3F). Gene set enrichment analysis revealed this response to be driven by enhanced type I interferon signaling (Fig 3G-H, S3B). This was validated at the protein level by elevated STAT1 phosphorylation (Fig S3C).

### Tumor PD-L1 loss drives transcriptional changes in myeloid cells, not CTLs, in the metastatic niche

We then sought to explore how tPD-L1 suppressed type I interferon signaling. We did so through single-cell RNA sequencing of CD45^+^ leukocytes from 4T1 WT and PDL1-KO colonized lungs (Fig 4A). Following shared nearest neighbor clustering and dimensional reduction of ~7500 cells by UMAP, cell populations were identified by comparison to publicly available datasets (Fig. 4B). We examined the expression of genes that were found to be differentially expressed in our bulk RNA-experiment in each cell population at the single-cell level. Surprisingly, analysis of differentially expressed genes showed that expression changes resulted from increased transcription by macrophages (Fig 4C). Similarly, application of a signature composed of genes repressed by tPD-L1 in bulk RNA sequencing data (referred to as tPD-L1 signature) showed the greatest magnitude change centered on macrophages (Fig 4D).

**Figure 4.**
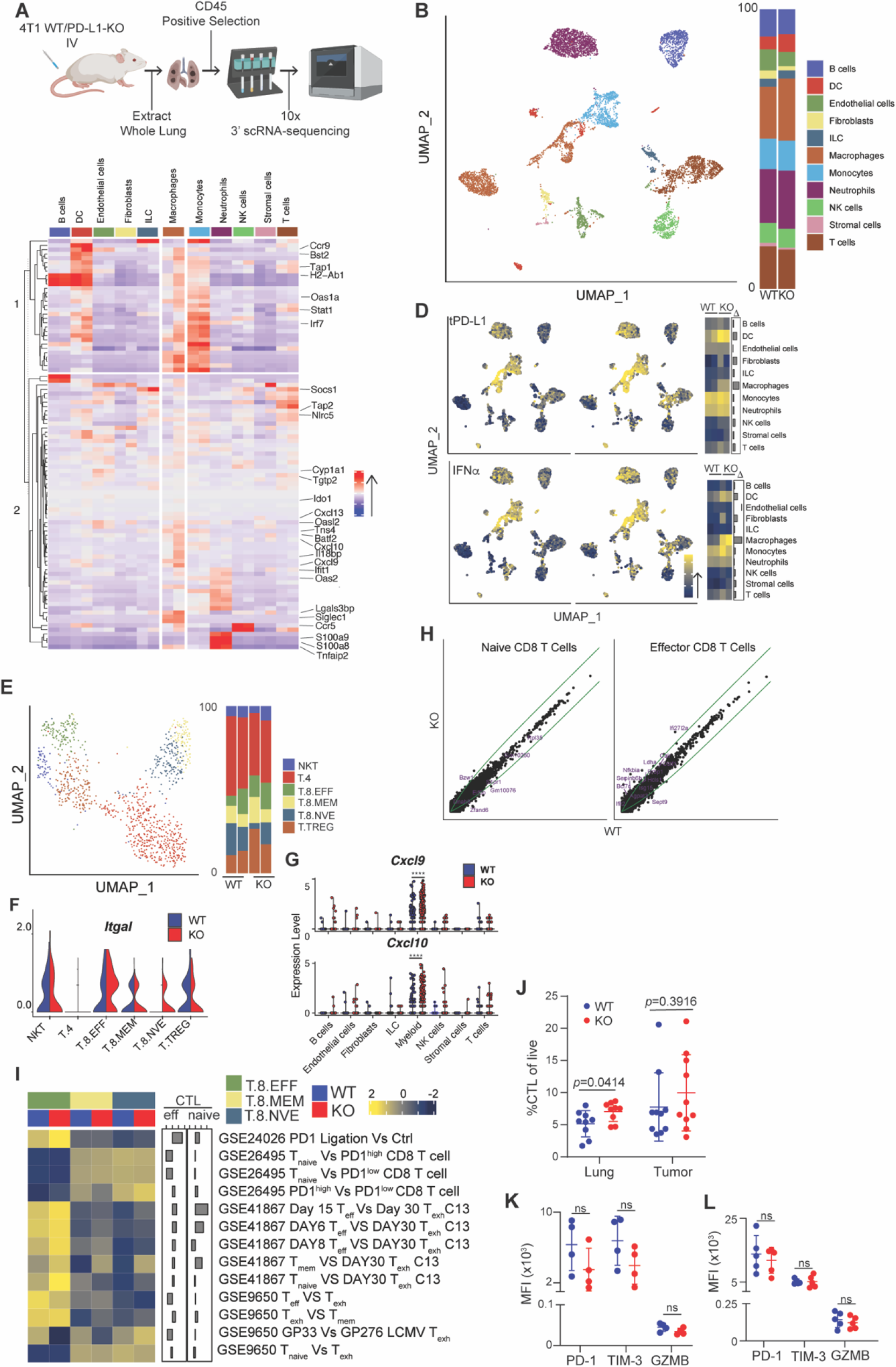
CTL activation and differentiation is independent of tPD-L1. **A)** Schematic for scRNA-sequencing of 4T1-colonized lungs. **B)** UMAP projection (left) and barplot of identities of CD45^+^ cells (right) in 4T1 colonized lungs. **C)** Heatmap of indicated transcript expression in listed cell populations. For each cell population, left column represents WT expression, while right column represents KO expression. **D)** Single-cell scoring of tPD-L1 loss (top) and IFNα (bottom) signature. Left plot includes cells derived from WT-colonized lungs, while right column represents KO-colonized lungs. **E)** UMAP projection of k-means clusters of T Cells. F-G) Violen plots of indicated transcript. NKT: NK T cells, T.4: CD4 T cells, T.8.EFF: Effector CD8 T cells, T.8.MEM: Memory CD8 T cells, T.8.NVE: Naïve CD8 T cells, T.TREG: Treg cells **H)** Expression of all transcripts in WT (x-axis) and KO (y-axis) T cells of indicated populations. Green lines represent 1.5 fold difference in transcription between populations. **I)** Expression of exhaustion-associated signatures in each cluster. **J)** CTL infiltration of primary tumor or lung metastases. N=10/condition **K-L)** Surface marker expression of CTLs isolated from lung metastases (K) or primary tumor (L). N=5/condition. All error bars in figure represent mean ± SD. **** p < 1*10^−4^

Contrastingly, tPD-L1 had minimal impact on the CTL transcriptome. Sub-clustering of the T cell compartment showed no changes in the relative proportions of activated versus naïve CTLs (Fig 4E). However, markedly enhanced expression of the integrin *Itgal* (encodes LFA-1), which mediates firm adhesion to vascular endothelium ^21^, provides a rationale for increased CTL infiltration as detected on IHC (Fig 4F). In addition to the increased homing-related gene, loss of tPD-L1 resulted in dramatic increases in myeloid expression of the chemokines *Cxcl9* and *Cxcl10*, which are ligands for CXCR3 that promotes T cell tumor infiltration (Fig. 4G). To examine whether tPD-L1 drove T cell exhaustion in the experimental metastasis model, we scored each T cell subset against a variety of transcriptional signatures associated with exhaustion or PD1-mediated dysfunction. No consistent difference was observed in signature scores or transcriptomes, suggesting that, in the metastatic setting, tPD-L1 does not drive CTL exhaustion (Fig 4H-I). Consistent with our histologic and transcriptomic data, flow cytometric analysis showed selectively enhanced infiltration of CTLs in lungs colonized with tPD-L1 deficient 4T1 (Fig 4J). We then sought to validate our transcriptome-level data in CTLs at the protein level. No difference was observed in the expression of exhaustion-(PD-1, TIM-3) or effector-(Granzyme B) associated markers by CTLs infiltrating the primary tumor or lung-resident metastases (Fig 4K-L, S4A & B). Together, these findings demonstrate that tPD-L1 restrains the recruitment, not the functionality, of CTLs.

As myeloid cells show the greatest transcriptional change after PD-L1 loss, we then investigated the myeloid compartment. To analyze the differentiation trajectory of myeloid cells, we performed PHATE analysis ^22^ followed by k-means clustering, which reveals a central progenitor population (cluster MA) that diverges into two branches based on low (cluster MB) and high (cluster MC) type I interferon signaling (Fig 5A-B, S5A-B). tPD-L1 antagonizes myeloid cell differentiation into cluster MC, reducing the expression of antigen presentation machinery (*B2m*, *H2-DMb1*) and T cell-recruiting chemokines (*Cxcl9*, *Cxcl10*) (Fig 5C, S5C). The alternative cellular fate adopted by tPD-L1 resident macrophages correlated with enhanced myeloid SHP-2 signaling, a downstream effector of PD-1, and decreased type I interferon, indicating that tPD-L1 may directly restrict myeloid differentiation by engaging mPD-1 (Fig 5D).

**Figure 5.**
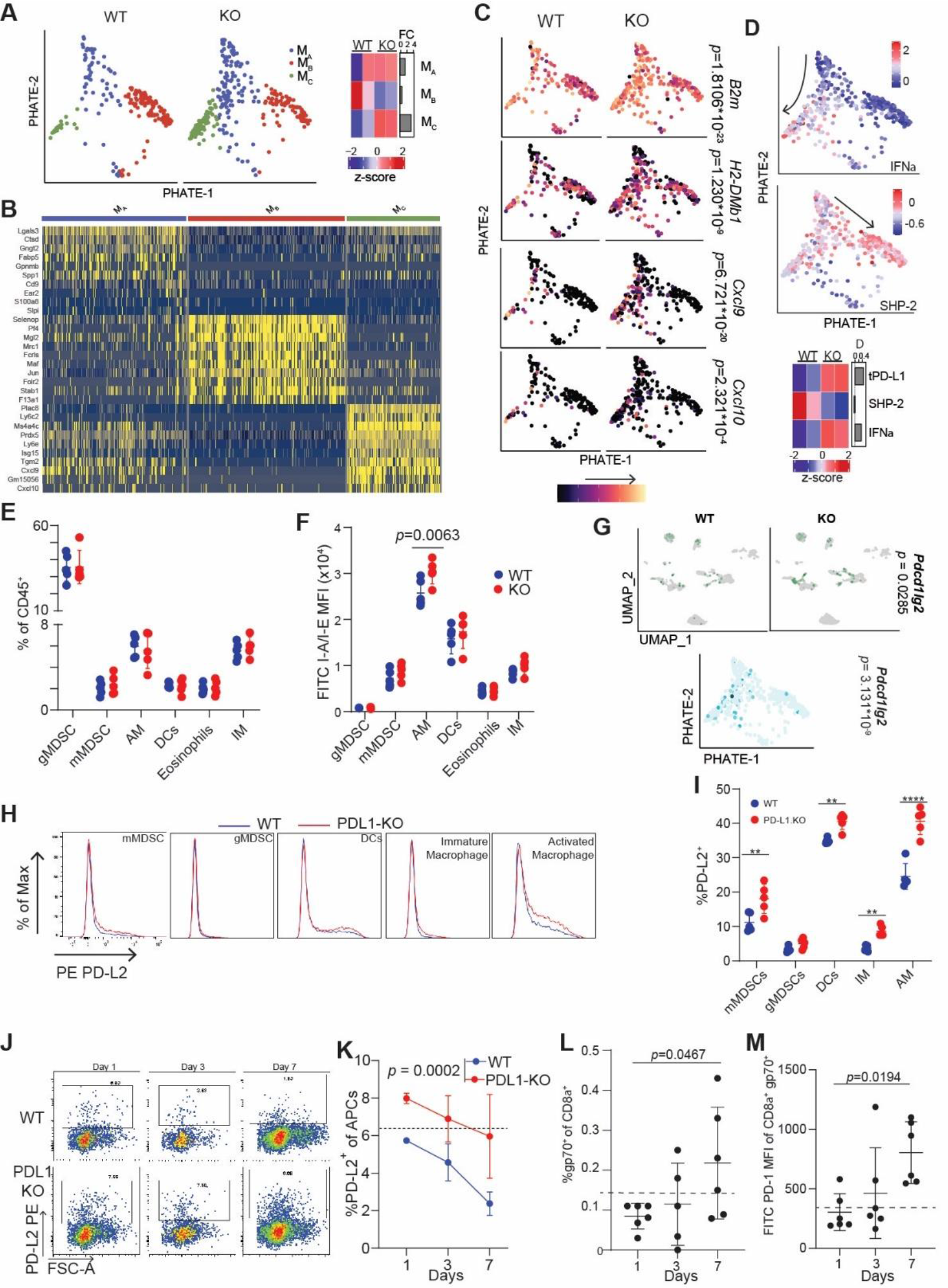
tPD-L1 restrains type I interferon-driven myeloid cell activation. **A)** PHATE projection of non-alveolar macrophage population (left) and proportion of cells belonging to each cluster (right). **B)** Dimensional heatmap of top ten most variably expressed genes in each cluster. **C-D)** Expression of indicated genes (C) and signatures (D) in cells. P-value calculated by Seurat “FindMarkers” function of cluster M_C_ vs M_B_ expression. **E-F)** Myeloid composition (E) and MHC class II expression (F) fourteen days post-colonization. Two independent experiments. N = 3,5/condition. Two-way ANOVA with Sidak’s multiple comparison. **G)** Differential expression of PD-L2 in lungs colonized by 4T1 WT or PD-L1-KO. **H**) Representative histograms of PD-L2 surface expression on indicated cell populations after colonization by 4T1 WT (blue) or 4T1 PDL1-KO (red). **I)** Quantification of %PD-L2^+^ cells in indicated cell populations. N=5/condition. **J-K)** Representative plots (J) and quantification (K) of %PD-L2^+^ antigen-presenting cells (CD45^+^CD11c^+^Ly6G^−^) at indicated time points. Results are representative of two independent experiments. Two-way ANOVA. N=3,5/condition. **L-M)** Quantification (L) and PD-1 expression (M) of gp70-tetramer^+^ cells in 4T1 WT or PDL1-KO colonized lungs at indicated timepoints. N=6/condition. Dotted line represents FMO baseline.Results are representative of two independent experiments. Fisher’s Exact Test. All represent mean ± SD. ** p < 0.01; **** p < 0.0001

To confirm these transcript-level observations, we performed flow cytometric analysis of 4T1-colonized lungs. Validating our transcriptomic data, no change in bulk lung myeloid cell composition was observed, though enhanced antigen presentation machinery was detected in macrophages (Fig 5E-F). Given that type I interferon signals in an autocrine or distance-dependent paracrine manner ^23^, we sought to confirm that tPD-L1 was directly acting on myeloid cells to suppress type I interferon production and signaling. Transcriptomic and proteomic analysis identified *Pdcd1lg2*, which encodes PD-L2, as a marker for both type I interferon-stimulated myeloid cells after tPD-L1 loss (Figure 5G-I, S5D-F). Differential expression of PD-L2 was observed in 4T1-colonized lungs as early as one day after injection (Figure 5J-K). This increase occurred before a robust infiltration and activation of 4T1-specific CTLs, further indicating remodeling of myeloid cell population occurred independently of CTL PD-1 (Fig. 5L-M, S5G-H).

### tPD-L1 engagement with mPD-1, not CTL PD-1, creates an immunosuppressive niche

Given previous reports of PD-1 expression on macrophages and dendritic cells ^14, 16^, as well as the enhanced SHP-2 signaling observed in tPD-L1 colonized lungs, we hypothesized that this attenuation of type I interferon signaling was mediated by tPD-L1 engagement with PD-1 found on lung-resident myeloid cells. Myeloid cell surface PD-1 expression was enhanced following tPD-L1 loss (Fig 6A-B). To assess the consequences of the tPD-L1:mPD-1 interaction, 4T1 cells were co-cultured in vitro with RAW264.7 macrophages. RAW surface expression of PD-1 was upregulated in a cell-contact and IFNAR-dependent manner following co-culture with 4T1 (Fig 6C, S6A-E). Engagement with 4T1 tPD-L1 enhanced SHP-2 activation and decreased pSTAT1 levels in RAW cells, diminishing expression of MHC class II both on 4T1 and RAW cells (Fig. 6D-F). This phenomenon was confirmed in vivo, as flow cytometry also revealed elevated MHCII expression on CD45^neg^ stromal and tumor cells following loss of tPD-L1 (Fig 6G). Similarly, RAW264.7 PD-1 loss led to increased sensitivity to stimulation by the TLR3 ligand poly(I:C) both on RAW cells and 4T1 in a co-culture system, as measured by the surface marker BST2 (Fig 6H-I, Fig S6F-G). We then hypothesized that RAW PD-1 engagement suppressed autocrine and paracrine type I interferon release. Accordingly, disruption of RAW PD-1 expression or 4T1 PD-L1 expression enhanced IFNβ production (Fig. 6J). Given the identity of IFNα as a potent PD-L1/L2 inducer ^24^, these findings demonstrate the existence of a negative feedback circuit in which the PD-1/L1 axis is induced by and then attenuates type I interferon signaling.

**Figure 6.**
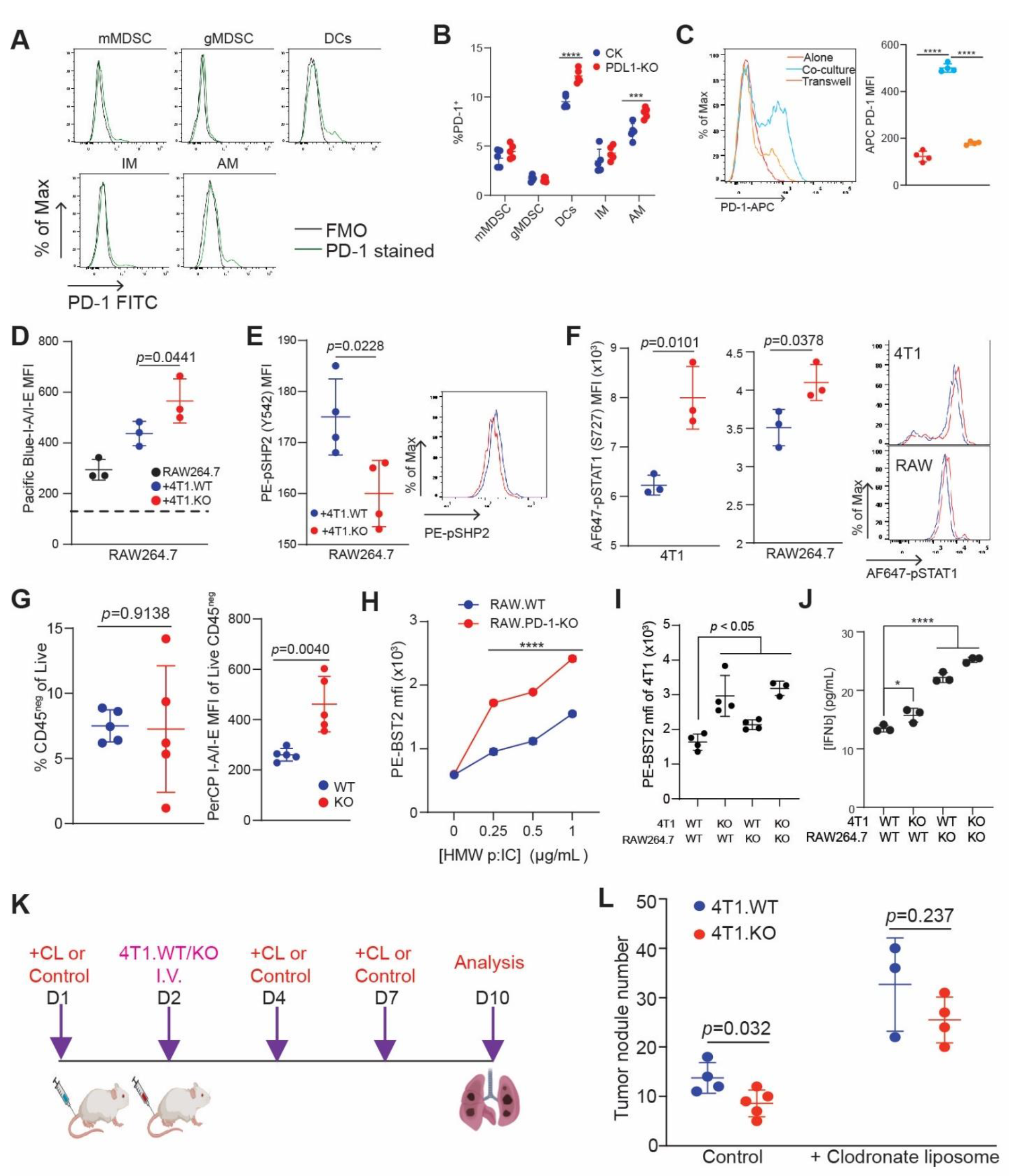
Myeloid PD-1 connects tumor PD-L1 to CTL suppression through suppressing antigen-presentation and interferon signaling following tPD-L1 engagement. **A-B)** Representative histograms (A) and quantification (B) of PD-1 expression on indicated cell populations isolated from 4T1 WT or PDL1-KO colonized lungs fourteen days after injection. N=5/condition. Two independent experiments. **C)** RAW264.7 expression of PD-1 following overnight co-culture with 4T1 cells with and without 4T1-RAW264.7 cell-cell contact. N=3/condition. Two independent experiments. One-way ANOVA with Dunnett’s multiple comparison. **D-F)** MHC class II (D), pSHP2 (Y580) (E), and pSTAT1(S727) (F) expression on RAW (D-F) and 4T1 (F) cells after co-culture. Two independent experiments. N=3-4/condition. **G)** CD45^neg^ percentage composition (left) and MHC class II expression (right) seven days post-colonization. Two independent experiments. N = 3,5/condition. **H-I)** Expression of BST2 following three days stimulation of indicated cell lines with HMW poly(I:C) (H) or co-culture of indicated cell lines (I). N=4/condition. Two-way ANOVA, Sidak’s test **J)** IFNβ levels in co-culture following 24 hours of stimulation with HMW poly(I:C). Dunnett’s multiple comparison. All represent mean ± SD. ****p<0.001; ****p<0.0001. **K**. Myeloid cell depletion scheme. WT and PD-L1 KO 4T1 tumor-bearing mice were treated with liposome control and clodronate liposome, respectively one day prior tumor cell injection. The tumor-bearing mice were treated every 3 days with liposome control or clodronate liposome for 3 times. **L**. Mouse lungs were inflated with India ink and quantified for tumor nodule numbers.

Our above findings determined that mPD-1 and its downstream signaling pathway is the target of tPD-L1. To functionally demonstrate that myeloid cells bridge tPD-L1 expression to CTL suppression, we depleted myeloid cells by clodronate liposomes in tumor-bearing mice and analyzed tumor growth in the lungs. Depletion of myeloid cells diminished tPD-L1 function in promoting tumor growth in the lungs in this experimental lung metastasis model (Fig. 6K-L). We therefore concluded that mPD-1 connects tPD-L1 to CTL suppression and thus promotes lung metastasis.

### Autocrine type I interferon signaling in myeloid cells drives CTL recruitment in the metastatic tumor and response to checkpoint blockade immunotherapy in human cancer patients

To determine the human relevance of our findings, we extended our above findings to a cohort of human patients with metastatic basal cell carcinoma who had single cell sequencing performed before and after PD-1 blockade ^25^. Myeloid cell PHATE analysis revealed differentiation restriction that was released following PD-1 blockade, leading to enhanced interferon signaling, antigen presentation, and CTL-recruiting chemokine expression (Fig 7A-C). Given the transcriptional heterogeneity between mouse and human myeloid cells, we transferred mouse myeloid cell cluster identities to a human lung myeloid cell scRNAseq dataset to develop transcriptomic signatures of cluster-identifying genes ^26^. The MC signature was validated by correlation analysis across the TCGA database that demonstrated an inverse relationship between the MC gene signature and myeloid SHP-2 activity in all tumor types represented (Fig S7A).

**Figure 7.**
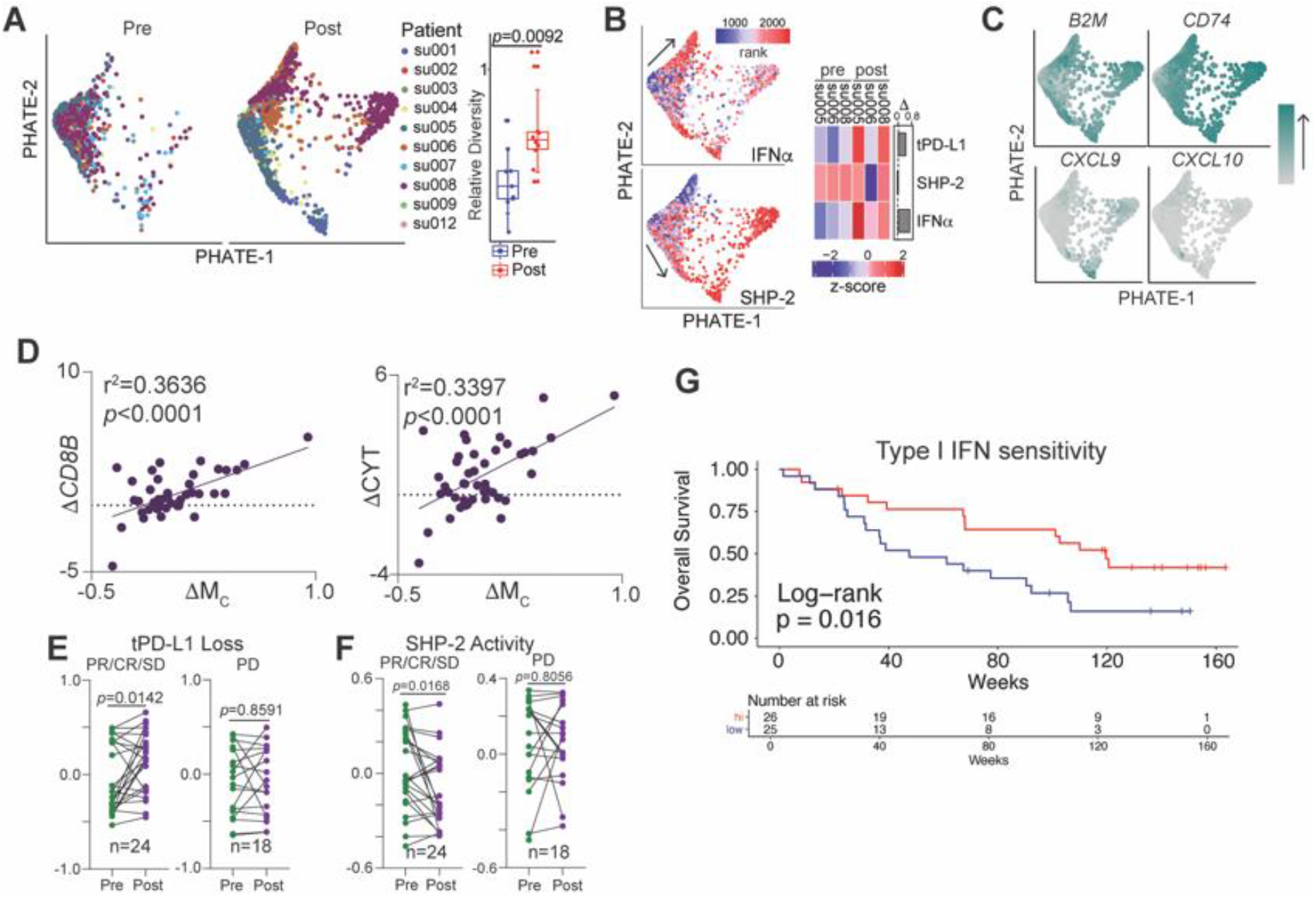
PD-1 blockade releases macrophage interferon signaling and production in human cancer patients. **A)** PHATE projection of metastatic basal cell cancer pre-and post-PD-1 blockade (left) and average cell displacement (see methods) from mean cell location, normalized per treatment (right). Student’s t test. **B)** Single cell scoring of indicated signatures (left) and heatmap of sample averages (right). **C)** Single cell expression of indicated genes, with expression imputed by Rmagic. **D)** Correlation of change in indicated transcripts after nivolumab treatment in metastatic melanoma cohort (BMS038). **E-F)** Change in tPD-L1 loss and SHP2 activity signature (BMS038). Paired t-test. **F)** Correlation between change in indicated signatures pre- and post-nivolumab therapy (BMS038). **G)** Survival analysis of type I interferon responsive and nonresponsive tumors pre and post PD-1 blockade. Two-sided Log-rank survival test. (BMS038).

Analysis of a pre- and post-treatment RNA sequencing dataset of metastatic melanoma patients treated with nivolumab revealed positive associations between the MC gene signature and markers of CTL infiltration (*CD8B*) and cytotoxic activity (Fig. 7D). Consistent with what was observed in the metastatic mouse tumor models, the expression levels of CXCL9 and CXCL10 are higher post PD-1 blockade immunotherapy in the metastatic human tumor (Fig. 7C). These findings suggest that this myeloid cell population similarly drive CTL recruitment in humans (Fig 7D). This correlation was validated across all tumor types represented in the TCGA database (Fig S7A-C). Resembling our findings in the 4T1 model, however, when samples were normalized to CD8B levels to correct for CTL infiltration levels, the correlation was no longer significant (Fig S7D). This indicates that tumor-expressed PD-L1 may repress CTL recruitment, rather than directly modulating CTL activation in human cancer patients with metastatic disease.

Enhanced expression of the tPD-L1 loss signature and repression of myeloid SHP-2 activity was observed exclusively in responders (Fig 7E-F). Based this, we hypothesized that tumors that possess infiltration by SHP-2 inhibited myeloid cell progenitors (high MA, *PTPN11*) and rich capacity for interferon signaling (high *IFNAR1/2*) would perform superiorly following checkpoint blockade. Indeed, these patients demonstrated enhanced overall survival following checkpoint blockade (Fig 7G). Survival benefit was observed at later, not initial, time points, indicating that these factors impact the emergence of metastatic growths and are required for a durable response. Taken together, these findings indicate that the myeloid PD-1/SHP2/IFN-I/CXCL9&10 axis controls tPD-L1-mediated suppression of CTL tumor infiltration in human cancer patients with metastatic disease.

## Discussion

Work in the past decade has increasingly highlighted the role of PD-1 in non-canonical immune cell populations, mainly ILCs, dendritic cells, and on tumor cells themselves, in creating an immunosuppressive niche ^27, 28, 29^. Myeloid cells, in particularly, have long been noted to contribute to the formation of an immunosuppressive metastatic niche ^30, 31^. These pro-metastatic functions have historically been attributed to alternatively activated or M2 suppressive macrophage populations ^32^. Macrophage PD-1 has been shown to promote tumor immune evasion ^33^. However, the mechanisms controlling the phenotypic switch between suppressive and inflammatory, tumoricidal populations have remained unclear. In this study, we demonstrated that tPD-L1 exhibits no direct role in suppression of CTL activation and cytotoxicity when only the target tumor cells and tumor-specific CTLs are present in the microenvironment. Instead, we show that tumors previously considered to be resistant to PD-1/L1 blockade nonetheless depend on tPD-L1 expression to prevent immune clearance of metastases. In doing so, we uncouple the role of tPD-L1 at the metastatic site, where it binds to mPD-1 to enforce an immunologically cold microenvironment.

Conflicting findings for the role of tPD-L1 in contribution to PD-1 blockade efficacy have been reported, but there is a consensus that tPD-L1 may be sufficient, but not necessary, for response to PD-1 blockade in primary tumors ^34, 35, 36^. Our results indicate that tPD-L1 may control CTL recruitment versus functionality, supported by recent observations of simultaneous infiltration of metastases by CTLs following PD-1 blockade ^37^. Accordingly, recent publications have highlighted the importance of dendritic cell expressed PD-L1 in controlling T cell function, suggesting the effect of PD-1 blockade on CTLs may occur in secondary and tertiary lymphoid structures, rather than at the tumor site ^25, 38, 39^. We have thus demonstrated here that tPD-L1 has an independent role in metastasis.

IFNα was the first FDA-approved cancer immunotherapy for adjuvant treatment in melanoma and renal cell carcinoma, and, while its response rate was minimal, it induced long-term, metastasis-free remission in a significant fraction of patients with late-stage metastatic disease, similar to checkpoint blockade ^40, 41^. Our work and others have shown that type I interferon induces both PD-L1 and PD-1 in myeloid cells, though PD-L1 expression is up-regulated more rapidly ^42^. This supports the hypothesis that PD-1/L1 axis functions as a negative feedback circuit for type I interferon signaling and highlights the intriguing possibility that low response rates in interferon alpha therapies were due to compensatory PD-1/L1 upregulation. Supporting this, phase I/II clinical trials have shown early success for direct administration of interferon, or indirect upregulation through TLR and STING agonists, in combination with anti-PD-1/L1 ^43, 44^. Furthermore, forced overexpression of *Irf7*, which mediates the second wave of interferon production in response to type I interferon exposure, abrogated triple negative breast cancer bone metastasis and was silenced at spontaneously occurring metastasis, including at the lung ^45, 46^. Similarly, prostate cancer metastases display a silencing of type I interferon production ^47^. Interestingly, mutations in the type I and II interferon pathway (*JAK1*/*JAK2*, etc.) are associated with adaptive resistance to immunotherapy, suggesting that suppression of interferon signaling is a primary mechanism by which tumors achieve successful immune evasion and adaptive resistance ^48, 49^. In this study, we determine that type I interferon signaling pathway is essential for myeloid cell function in CTL recruitment to the lung metastasis niche. Mechanistically, we determine that the tPD-L1/mPD-1/IFN-I/STAT1/Cxcl9/10 axis controls CTL tumor infiltration in lung metastasis to promote lung metastasis, which underlies the efficacy of ICB immunotherapy in the neoadjuvant and the adjuvant setting ^8, 9, 10, 11, 12, 13^.

One limitation in this study is that we exclusively analyzed lung metastases, which due to their location in barrier tissue, may have a uniquely inflamed microenvironment ^50^. Further investigation is needed to analyze whether this mechanism operates analogously at non-lung metastatic sites. Metastasis accounts for over 90% mortality in human cancers. Such studies may elucidate the molecular mechanism underlying mPD1/IFN-I pathway in immune suppression and tumor metastasis in other metastatic settings and are thus of high significance in human ICB immunotherapy.

## Methods

### Mice

Mice were purchased from Jackson Laboratory (Bar Harbor, ME) or Charles River (Frederick, MD). All animal studies were approved in advance by Augusta University Institutional Animal Care and Use Committee (Protocol #: 20080162).

### Cell lines

CT26, 4T1, RAW264.7 cell lines were obtained from American Type Culture Collection (ATCC) (Manassas, VA). J774M cells were generated as previously described ^51^. 293FT cells were obtained from (ThermoFisher Scientific). CMS4-met cells were as previously described ^52^. AH1 antigen-specific T cells (2/20 CTLs) were generated and maintained as previously described ^53^. Cell lines were tested bi-monthly for mycoplasma and were mycoplasma-free at time of experiments.

### Gene knockout cell line generation

HEK293FT cells were co-transfected with pCMV-VSV-G (Addgene #8454), psPAX2 (Addgene #12260) and lentiCRISPRv2 (Genscript, Piscataway, NJ) plasmids using Lipofectamine 2000 (Life Technologies) to produce CRISPR lentivirus ^54^. sgRNA sequences can be found in Table S2. Tumor cells were transduced with the lentivirus particle. Cells were selected with puromycin and bulk cells were then sorted for marker**--** cells using a BD FACSARIA (BD Biosciences).

### Tumor-CTL Co-culture system

Tumor cells were cultured with 2/20 CTLs in a 96-well U-bottom plate. Floating and adherent cells were collected, stained, and analyzed by flow cytometry. To assay PD-L1-mediated inhibition of 2/20, 96 flat-bottom plates were coated with 1 μg/mL 145-2C11 (BioXcell), IgG (MOPC-21, BioXcell), or mPD-L1-FC (Biolegend, Cat#758208) and incubated at 4°C overnight before seeding cells.

### In vivo tumor models

4T1 orthotopic tumors were established by injection at the right second mammary gland. CT26 subcutaneous tumors were established by injection at the right flank. For experimental metastasis models, tumor cells were injected into the lateral tail vein. To analyze lung metastasis, lungs were harvested, inflated with an India ink solution, and quantified as previously described ^55^. For neutralization experiments, 200 μg of anti-CD8α (53-6.7, BioXcell) or isotype control (2A3, BioXcell) were injected i.v. every other day during the course of the experiment.

### Flow cytometry

Samples were blocked in FACS Buffer (PBS + 2%FBS) containing mouse FcBlock. Antibodies and other reagents were listed in Table S1. For co-culture cytotoxicity experiments, supernatant and cells were harvested and stained with APC-AnnexinV and propidium iodide (PI). For intracellular staining, samples were stained for surface antigens and viability, then cells were fixed and permeabilized using Cytofix/Cytoperm Plus kit according to manufacturer’s instructions (BD Biosciences). Samples were acquired on an FACSCalibur with CellQuestPro or LSRFortessa with BD Diva 8.01 (BD Biosciences). All flow cytometry data analysis was conducted with FlowJo v10.6.0 (BD Biosciences).

### Statistical analysis

Statistical analysis was conducted using Prism8 (Graphpad) and p-values were calculated by a two-tailed Student’s t-test. Significance between survival groups were computed by two-sided log-rank test (Survival package, R).

### Bulk RNA Sequencing

Mice were injected i.v. through the lateral tail vein with 4T1.WT or 4T1.PDL1-KO cells. After seven days, mice were sacrificed, lungs were perfused with PBS, and mechanically homogenized and RNA was extracted using Direct-zol RNA microprep kit according to manufacturer’s instructions (Zymo Research, Irvine CA). RNA was sequenced by Novogene. Reference genome (mm10) and gene model annotation files were downloaded from Ensembl directly. Indices were built and paired-end clean reads were aligned to the reference genome using STARv2.5. Reads were quantified using HTSeqv0.6.1. Rank-log transformed normalized counts from DESeq2 were used as inputs for GSEA, GO and PCA analysis. GSEA was performed using GSEA4.0 (Broad) with gene-set permutation. GO pathway enrichment was performed with clusterprofiler.

### Single-Cell RNA Sequencing

Balb/c mice were injected i.v. with 4T1.WT or 4T1.PDL1-KO cells. After fourteen days, mice were sacrificed and lungs were perfused with PBS and harvested. Following collagenase digestion, leukocytes were positively selected using CD45 magnetic nanobeads (Biolegend). Cell viability was assessed using trypan blue and assured of ≥ 80% viable cells. Cells were loaded at a concentration to capture approximately 2-3 × 10^3^ targeted cells on a Chromium Chip B (10x Genomics). scRNA-seq libraries were generated using the Chromium Single Cell 3’ Reagent Kit v3 (10x Genomics) according to the manufacturer’s instructions. The libraries were sequenced on Illumina NextSeq 500 platform under the following sequencing protocol: 28 bp (Read 1), 8 bp (indexing Run), and 91 bp (RNA Read 2). Reads from the raw fastq files were mapped to mm10 Mouse Genome reference by STAR aligner in Cell Ranger 3.1.0 pipeline. Alignment generated 30-39K reads/cell with 75-102 million reads per sample at a target of 2.3-2.7 × 10^3^ cells per sample identified at 88% in Q30 Bases in RNA read as well as greater than 91% genome-mapping rates along with approximately 1600 median genes per cell. Filtered counts data was loaded into Seurat. High quality cells (200 < feature count <4000, percent mitochondrial reads < 10) were extracted for downstream analysis. Doublet clusters were manually identified and removed, then cells were subjected to doublet removal by DoubletFinder. Cells were then reclustered and remaining doublet clusters were again manually identified and removed based on the expression of lineage-defining genes. Cells were clustered using the first fifty principal components, and clusters were annotated by comparison to pre-existing RNA-sequencing datasets using SingleR. Myeloid and T cells clusters were then subsetted and reclustered based off of top twenty principal components for further analysis. MSigDB and signature sets generated during the course of this study were loaded into VISION for single-cell scoring.

### Signature Generation

tPD-L1 loss gene signature was created by isolating genes whose expression increased by at least two-fold with a p_adj_ of less than 0.05 calculated by DESeq2 following tPD-L1 loss from our bulk RNA-sequencing dataset. SHP-2 activation signature was calculated by isolating genes whose expression respectively increased two-fold (p_adj_ <0.05) in myeloid cells following treatment with the SHP-2 inhibitor SHP099. Score generated was then inversed. MA, MB and MC gene signatures were taken by using the Seurat “TransferData” function to transfer labels from our macrophage PHATE clusters to a previously published dataset of human lung tumor-resident macrophages ^26^. Cluster-identifying signatures were calculated by the Seurat “FindAllMarkers” function. All signatures generated in the course of this study can be found in Table S3.

### Patient Dataset Analysis

mRNA and survival datasets were extracted from TCGA database. Survival analyses were performed using the R survival and survminer packages. Data from the BMS038 was accessed from authors’ processed files (https://github.com/riazn/bms038_analysis) and normalized counts were generated following author’s scripts, which were used for downstream analyses ^56^. Previously published dataset of response following PD-1 treatment (PRJEB23709) was accessed from EMBL and pseudo-aligned with kallisto using default settings. Counts were loaded into DESEQ2 for downstream analysis, which was conducted using rlog values. To analyze tumor-infiltrating macrophages in a melanoma cohort, single cells annotated by the authors to be macrophages were subsetted and analyzed using Seurat pipeline. Batch effects were corrected by Harmony. To assess intrasample variation, mean displacement was calculated. Changes in melanoma cells resistant or naïve to ICB were accessed through the author’s dataset ^57^. Counts were extracted and analyzed analogously.

### Colonization Assay

Balb/c mice were injected i.v. with 4T1.WT or 4T1.PDL1-KO cells. After twenty four hours, mice were sacrificed, lungs were perfused with PBS, collected, minced with scissors, then homogenized in Trizol using an electric homogenizer. RNA was isolated according to the manufacturer’s instructions, and used for PCR analysis.

### Western Blotting

Lungs were collected and homogenized using an electric homogenizer. Lysates were subjected to Western blotting analysis. Antibodies are listed in Table S1.

### Immunohistochemistry

Lung and tumor tissue was fixed in 10% formalin and processed into paraffin blocks and cut into sections. Sections were stained and analyzed as previously described ^58^. Images were aquired using LAS4.1 software (Leica). CD3^+^ cells were quantified by quPATH.

### Cell Proliferation Assay

Cells were seeded into 96-well flat-bottomed plates. Cell count was determined using AqueousOne reagent (Promega) following manufacturers instructions. Reported values were normalized to Day 0 readings.

### Cytokine Stimulation

Bone marrow was harvested and cultured After three days of in vitro culture, bone marrow, J774M and RAW264.7 cells were harvested and cultured with the indicated factors for 48 hours: TNF-α (50ng/mL), GM-CSF (20 ng/mL), LPS (1μg/mL), IFNα (50 ng/mL), IFNβ (50 ng/mL), IFNγ (50 ng/mL), HMW poly(I:C) (1 μg/mL), PMA (50 ng/mL) and Ionomycin (50 ng/mL), or IL-6 (50 ng/mL).

### ELISA

RAW264.7 and 4T1 were treated with 50 ng/mL IFNβ and 30 ng/mL IFNγ, respectively, for 72 hours. 1×10^5^ of each cell line was combined in a flat-bottom 96 well plate and treated with 1 μg/mL HMW poly(I:C). After 24 hours, supernatant was harvested and IFNβ levels were quantified by Lumikine mouse IFNβ luminescent ELISA (Invivogen, Cat# luex-mifnbv2) according to manufacturer’s instructions.

### Software

Rstudio v1.2.1335 was used as working environment for R 3.6.3. Following R packages were used in data analysis: viridis_0.5.1, viridisLite_0.3.0, DoubletFinder_2.0.2, Seurat_3.1.4, biomaRt_2.40.5, ComplexHeatmap_2.3.2, factoextra_1.0.6, FactoMineR_2.3, GSVA_1.34.0, EnhancedVolcano_1.2.0, DESeq2_1.24.0, tidyverse_1.3.0, survminer_0.4.6, ggpubr_0.2.5, Rmagic_2.0.3, survival_3.1-8, SingleR_1.0.6, SingleR VISION_2.1.0.

## Author Contributions

JDK, PSR, GZ, DHM, and KL conceived and designed the project. JDK, PSR, CL, ADS, DSB and DY performed experiments. JDK, PSR and KL analyzed data. GZ and DHM, critical review of the manuscript. JDK and KL wrote the manuscript.

## Acknowledgements

We thank Ms. Penny Roon and Donna Kumiski at the Medical College of Georgia Electron Microscopy and Histology Core for technical assistance in tissue preparation and histology work. CD3 immunohistochemical staining was performed by the University of Georgia Veterinary school histology core. Cell sorting was performed by Dr. Jeanene Pikhala at the Georgia Cancer Center Flow Cytometry Core. 10x single-cell data acquisition was performed by Sam Chang-Sheng, Ph.D. at the Bioinformatics shared resource affiliated with Integrated Genomic and High-Performance Computing core facility at Georgia Cancer Center where the sample preparation was conducted by Eiko Kitamura Ph.D. This work was supported by National Institutes of Health (R01CA133085 R01CA227433 to K.L., F30CA236436 to J.D.K.) and VA Merit Review Award (I01CX001364 to K.L.).

