## Supplemental Information for "Tumor PD-L1 selectively suppresses type I interferon in myeloid cells to suppress CTL recruitment to promote lung metastasis"

### Supplemental Figures

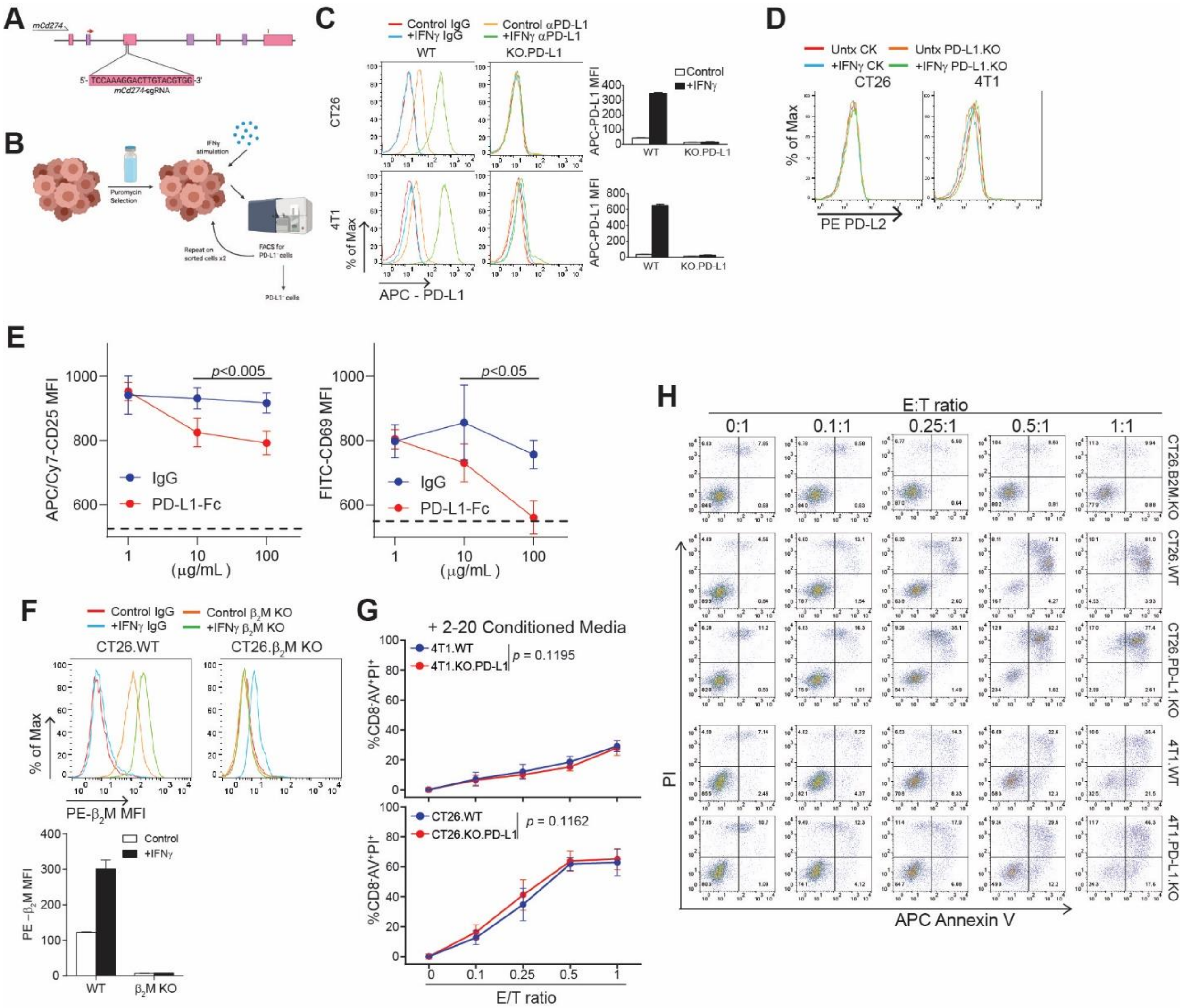

**Figure S1. Validation of co-culture system. A)** Schematic detailing design of sgRNA for targeted knockout of *Cd274*. **B)** Schematic detailing PDL1-KO cell line generation. Briefly, cells were transduced with lentiviral particles containing nontargeting sgRNA (WT) or *Cd274*-targeting sgRNA (PDL1-KO). Following puromycin selection, cells were stimulated with IFNγ overnight, then sorted for PD-L1 negative cells. Sorting was repeated twice. WT cell lines were analogously sorted once for PD-L1$^+$ cells. **C)** Validation of successful PD-L1 knockout in CT26 and 4T1 cell line. N=3/condition. Results are representative of two independent experiments **D)** Surface expression of PD-L2 on WT and PDL1-KO lines, with and without overnight treatment with IFNγ. **E)** Surface expression of activation markers on 2/20 CTL following overnight stimulation with 1 μg/mL αCD3ε and indicated concentrations of plate-bound IgG or PD-L1-Fc fusion protein. Dotted line represents MFI of unstimulated 2/20 CTL. Sidak's multiple comparison. N=3/condition. **F)** Validation of successful B2M knockout in CT26 cells. N=3/condition. Two-independent experiments. **G)** Cell death after overnight co-culture following no pretreatment with 2/20-conditioned media. N=6/condition. **H)** Representative images of WT and PDL1-KO tumor cell apoptosis following overnight co-culture with 2/20 CTLs at indicated ratios. Plots are gated on CD8α$^-$ cells. All represent mean ± SD.

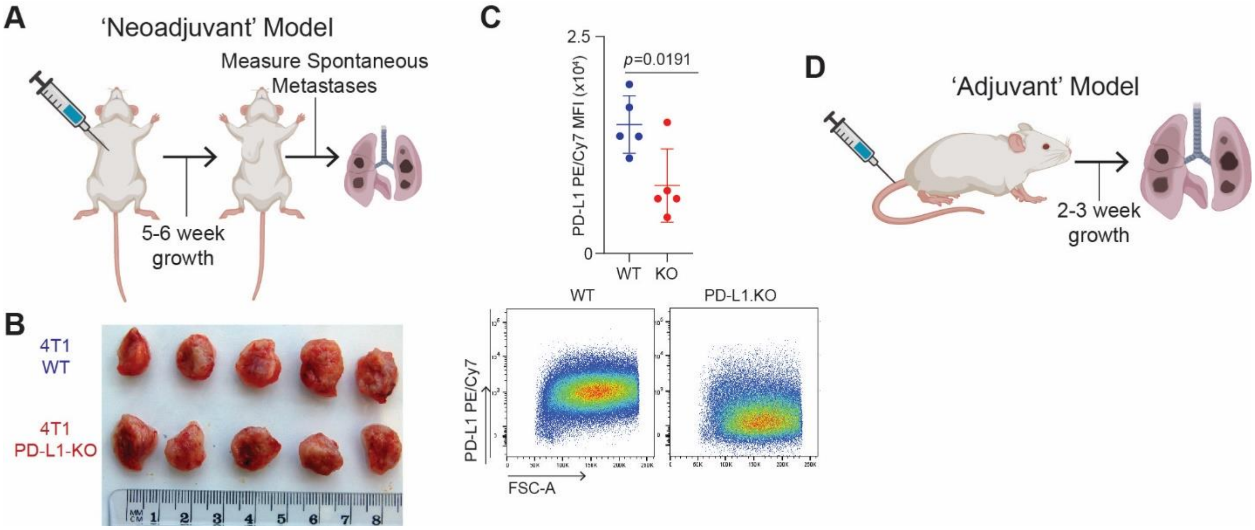

**Figure S2. In vivo metastasis models.**
**A)** Schematic detailing spontaneous metastasis model. **B)** Representative tumor growth of indicated cell lines following orthotopic injection, from Fig. 2A. **C)** Expression of tPD-L1 by 4T1 after orthotopic injection. N=5/condition. Three independent experiments. **D)** Schematic detailing spontaneous metastasis model.

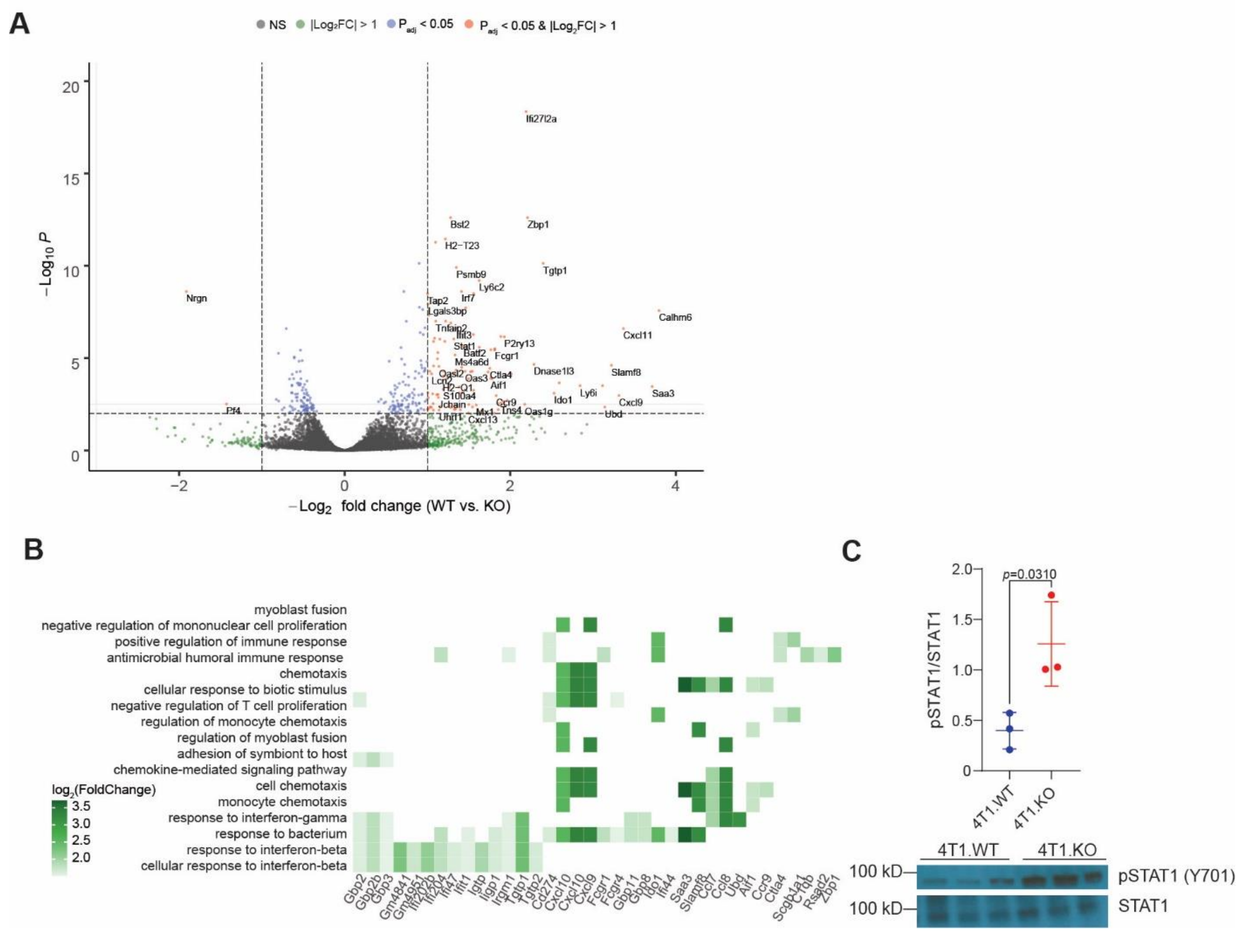

**Figure S3. tPD-L1 suppresses a Type I Interferon-driven response.**
A). Volcano plot of differentially expressed genes from bulk RNA-sequencing. B) Heatmap of GO terms enriched in PDL1-KO colonized lungs. C). Quantification (top) and Western blot (bottom) of pSTAT1(Y701) normalized to total STAT1 expression in 4T1.WT or 4T1.PDL1-KO colonized lungs seven days after experimental metastasis. Mean ± SD.

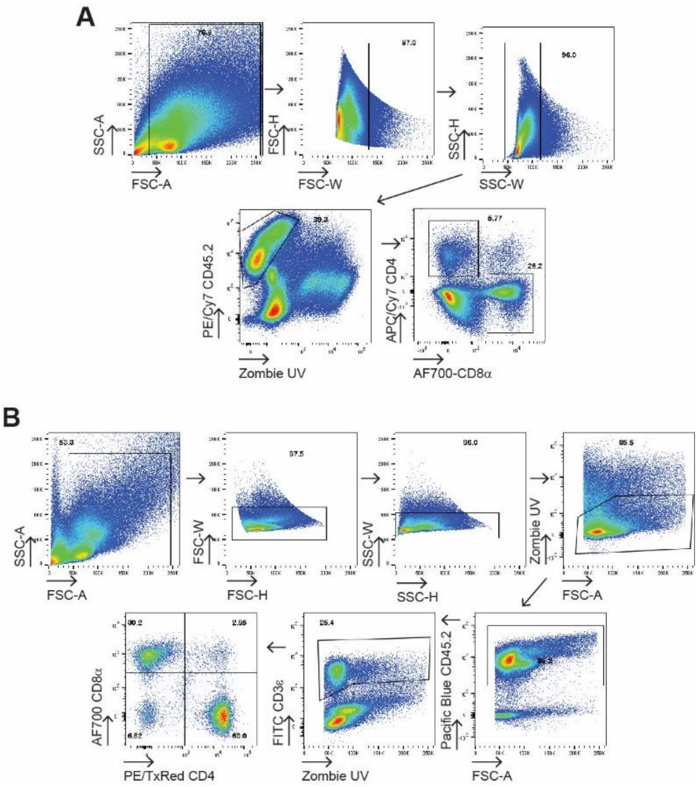

**Figure S4. Differential CTL infiltration in lung metastases and primary tumor.**
**A-B:** Representative gating for Tumor (A) and Lung(B) infiltrating T cells for Fig. 4I-K.

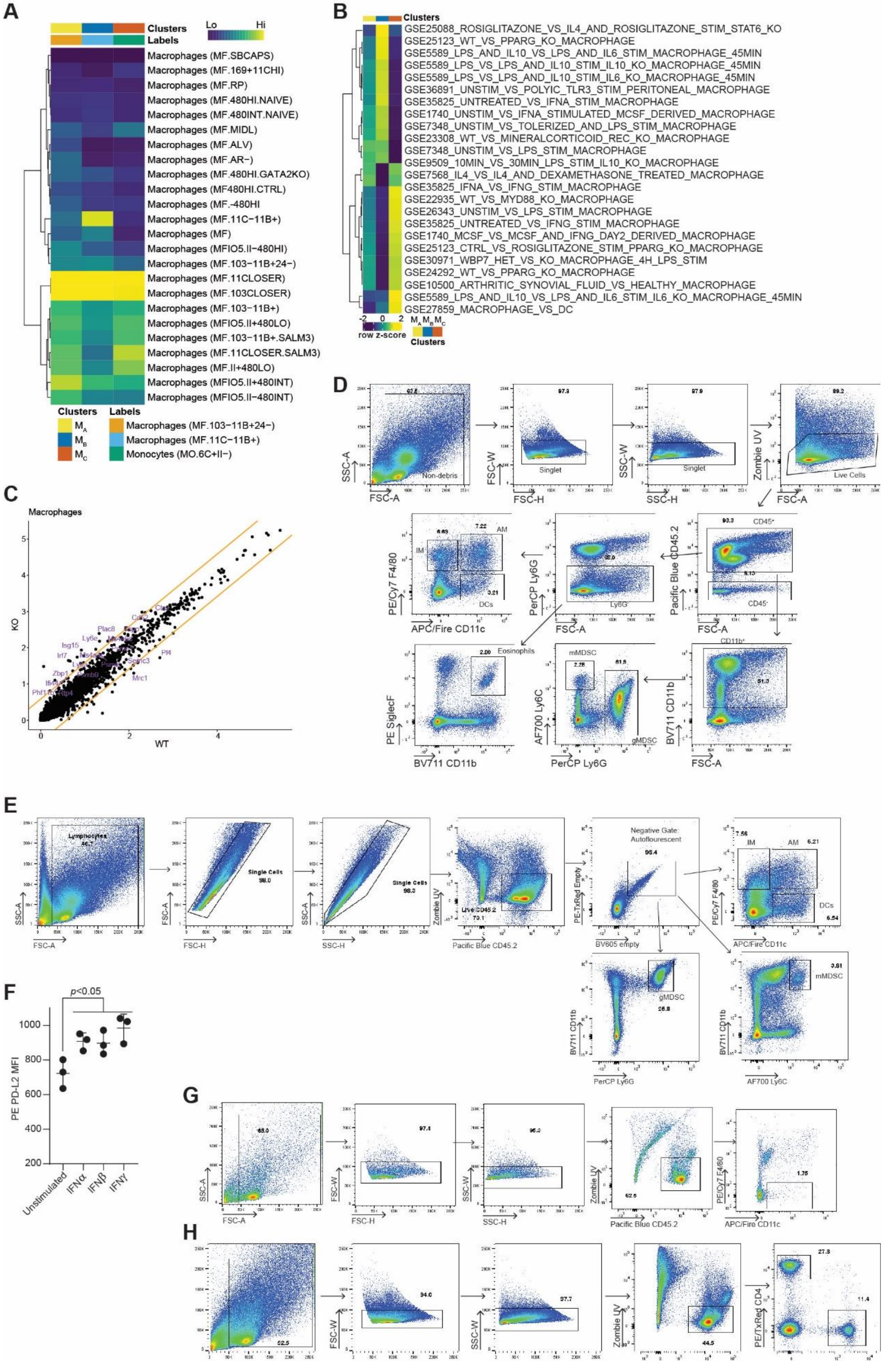
A
Clusters
Labels
Lo
Hi
Macrophages (MF.SBCAPS)
Macrophages (MF.169+11CHI)
Macrophages (MF.RP)
Macrophages (MF.480HI.NAIVE)
Macrophages (MF.480INT.NAIVE)
Macrophages (MF.MIDL)
Macrophages (MF.ALV)
Macrophages (MF.AR−)
Macrophages (MF.480HI.GATA2KO)
Macrophages (MF480HI.CTRL)
Macrophages (MF.-480HI)
Macrophages (MF.11C−11B+)
Macrophages (MF)
Macrophages (MFIO5.II−480HI)
Macrophages (MF.103−11B+24−)
Macrophages (MF.11CLOSER)
Macrophages (MF.103CLOSER)
Macrophages (MF.103−11B+)
Macrophages (MFIO5.II+480LO)
Macrophages (MF.103−11B+.SALM3)
Macrophages (MF.11CLOSER.SALM3)
Macrophages (MF.II+480LO)
Macrophages (MFIO5.II+480INT)
Macrophages (MFIO5.II−480INT)
Clusters
Labels
M_A
M_B
M_C
Macrophages (MF.103−11B+24−)
Macrophages (MF.11C−11B+)
Monocytes (MO.6C+II−)
B
Clusters
GSE25088_ROSIGLITAZONE_VS_IL4_AND_ROSIGLITAZONE_STIM_STAT6_KO
GSE25123_WT_VS_PPARG_KO_MACROPHAGE
GSE5589_LPS_AND_IL10_VS_LPS_AND_IL6_STIM_MACROPHAGE_45MIN
GSE5589_LPS_VS_LPS_AND_IL10_STIM_IL10_KO_MACROPHAGE_45MIN
GSE5589_LPS_VS_LPS_AND_IL10_STIM_IL6_KO_MACROPHAGE_45MIN
GSE36891_UNSTIM_VS_POLYIC_TLR3_STIM_PERITONEAL_MACROPHAGE
GSE35825_UNTREATED_VS_IFNA_STIM_MACROPHAGE
GSE1740_UNSTIM_VS_IFNA_STIMULATED_MCSF_DERIVED_MACROPHAGE
GSE7348_UNSTIM_VS_TOLERIZED_AND_LPS_STIM_MACROPHAGE
GSE23308_WT_VS_MINERALCORTICOID_REC_KO_MACROPHAGE
GSE7348_UNSTIM_VS_LPS_STIM_MACROPHAGE
GSE9509_10MIN_VS_30MIN_LPS_STIM_IL10_KO_MACROPHAGE
GSE7568_IL4_VS_IL4_AND_DEXAMETHASONE_TREATED_MACROPHAGE
GSE35825_IFNA_VS_IFNG_STIM_MACROPHAGE
GSE22935_WT_VS_MYD88_KO_MACROPHAGE
GSE26343_UNSTIM_VS_LPS_STIM_MACROPHAGE
GSE35825_UNTREATED_VS_IFNG_STIM_MACROPHAGE
GSE1740_MCSF_VS_MCSF_AND_IFNG_DAY2_DERIVED_MACROPHAGE
GSE25123_CTRL_VS_ROSIGLITAZONE_STIM_PPARG_KO_MACROPHAGE
GSE30971_WBP7_HET_VS_KO_MACROPHAGE_4H_LPS_STIM
GSE24292_WT_VS_PPARG_KO_MACROPHAGE
GSE10500_ARTHRITIC_SYNOVIAL_FLUID_VS_HEALTHY_MACROPHAGE
GSE5589_LPS_AND_IL10_VS_LPS_AND_IL6_STIM_IL6_KO_MACROPHAGE_45MIN
GSE27859_MACROPHAGE_VS_DC
-2 0 2
row z-score
M_A M_B M_C
Clusters
C
Macrophages
KO
WT
Isg15
Irf7
Ly6e
Plac8
Zbp1
Pf4
Mrc1
Sepnc3
D
SSC-A
FSC-A
Non-debris
FSC-W
FSC-H
Singlet
97.9
SSC-W
SSC-H
Singlet
97.9
Zombie UV
FSC-A
Live Cells
89.2
PE/Cy7 F4/80
APC/Fire CD11c
IM
AM
DCs
6.69
7.22
3.21
PerCP Ly6G
FSC-A
Ly6G⁻
Pacific Blue CD45.2
FSC-A
CD45⁺
CD45⁻
93.3
CD11b⁺
PE SiglecF
BV711 CD11b
Eosinophils
AF700 Ly6C
PerCP Ly6G
mMDSC
gMDSC
2.26
61.5
BV711 CD11b
FSC-A
E
Lymphocytes
SSC-A
FSC-A
Single Cells
98.0
FSC-A
FSC-H
Single Cells
SSC-A
SSC-H
Zombie UV
Pacific Blue CD45.2
Live CD45.2
Negative Gate:
Autoflourescent
PE-TxRed Empty
BV605 empty
96.4
IM
AM
7.56
6.21
PE/Cy7 F4/80
APC/Fire CD11c
DCs
6.54
BV711 CD11b
PerCP Ly6G
gMDSC
26.8
BV711 CD11b
AF700 Ly6C
mMDSC
3.81
F
p<0.05
PE PD-L2 MFI
1000
800
600
400
200
Unstimulated
IFNα
IFNβ
IFNγ
G
SSC-A
FSC-A
65.0
FSC-W
FSC-H
97.4
SSC-W
SSC-H
Zombie UV
Pacific Blue CD45.2
62.5
PE/Cy7 F4/80
APC/Fire CD11c
1.75
H
FSC-W
94.0
SSC-W
97.7
Zombie UV
44.5
PE/TxRed CD4
27.3
11.4

**Figure S5. Transcriptomic signature of Macrophage PHATE clusters and time-course of interferon-stimulated gene expression.**

**A)** SingleR comparison of PHATE clusters to Immgen transcriptional signatures. **B)** Heatmap of mean expression of single cell scores across indicated clusters. Signatures listed were those most variable across PHATE clusters. **C)** Differential gene expression in macrophages following tPD-L1 loss. Lines represent 2-fold change. **D)** Gating strategy for Fig. 5E-F. **E)** Gating Strategy for Fig 5H-I. **F)** Expression of PD-L2 by bone-marrow-derived myeloid cells after overnight stimulation with indicated cytokines. One-way ANOVA with Dunnett's. N=3/condition. Representative of two independent experiments. **G)** Gating strategy of Fig. 4J-K. **H)** Gating strategy of Fig. 5L.

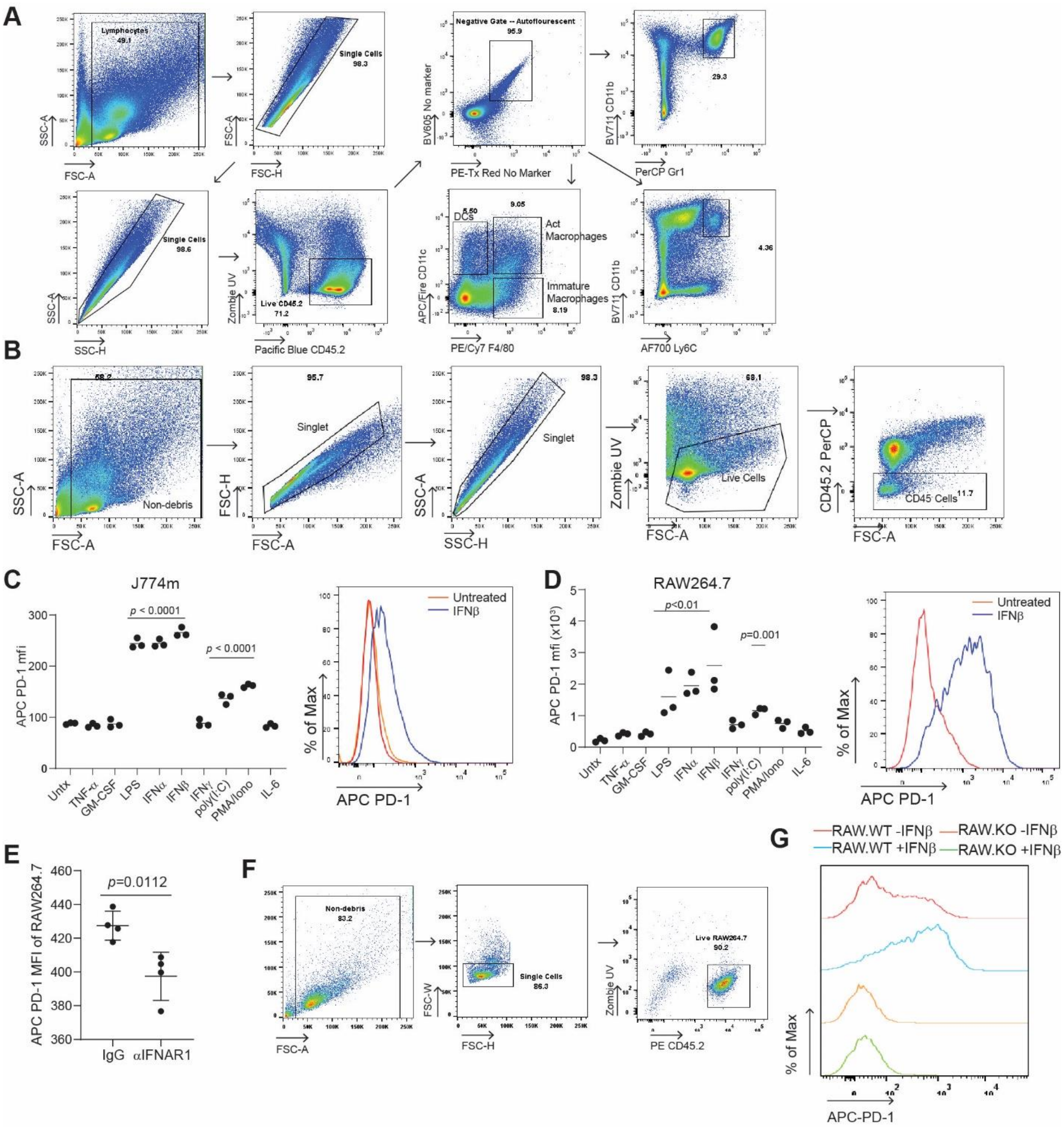

**Figure S6. Myeloid PD-1 is induced by Type I interferon.**

**A)** Representative gating strategy for Fig. 6A-B. **B)** Representative gating strategy for Fig. 6G. **C-D)** PD-1 expression in J774m (C) or RAW264.7 (D) cell lines following stimulation with indicated cell lines for 24 hours. Right: Representative histogram of PD-1 expression following IFNβ stimulation. Two independent experiments. N=3/condition. One-way ANOVA with Dunnett's multiple comparison. **E)** PD-1 expression (E in RAW264.7 following neutralization of IFNAR1 in co-culture with parent 4T1. N=4/condition. **F-G)** Gating strategy (F) and RAW264.7 WT and PD1-KO expression (G) of PD-1 72 hours post-stimulation with IFNβ. Histogram is representative of n=3/condition. Three independent experiments.

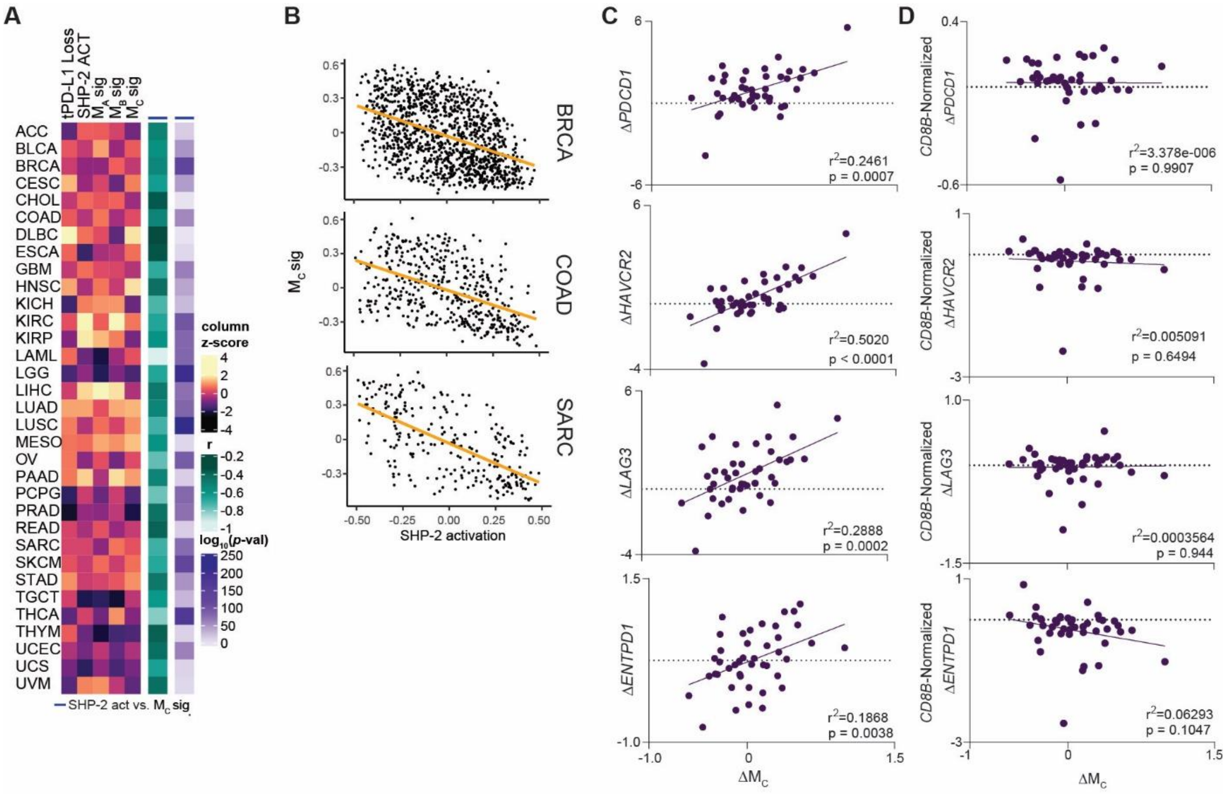

**Figure S7. SHP-2 activity antagonizes interferon-driven macrophage polarization across TCGA dataset.**

**A)** Relative expression of indicated signatures across TCGA dataset (left). Correlation and p-value between indicated scores (right). **B)** Representative correlation plots. **C and D)** Correlation between $M_C$ score and indicated markers of CTL activation/exhaustion before (C) and after (D) normalization to *CD8B* transcript level. p-values represent the probability that the slope is non-zero.

**Table S1. Antibodies**

| Antibody | Source | Identifier |
|---|---|---|
| APC-Annexin V (N/A) | Biolegend | 640941 |
| PE-Bst2 (eBio927) | eBiosciences | 12-3172-82; RRID:AB_763417 |
| APC-CD11b (M1/70) | BD Biosciences | 553312 |
| BV711-CD11b (M1/70) | Biolegend | 101242; RRID:AB_2563310 |
| APC/Fire-CD11c (N418) | Biolegend | 117352; RRID:AB_2572124 |
| BV785-CD11c (N418) | Biolegend | 117335; RRID:AB_11219204 |
| PE-CD16/32 (93) | Biolegend | 101308; RRID:AB_312807 |
| APC-CD25 (PC61) | Biolegend | 102012; RRID:AB_312861 |
| PE-CD3e (145-2C11) | Biolegend | 100308; RRID:AB_312673 |
| PE/Cy7-CD3e (145-2C11) | Biolegend | 100320; RRID:AB_312685 |
| PE/Dazzle-CD4 (RM4-5) | Biolegend | 100566; RRID:AB_2563684 |
| PE-CD45.2 (104) | Biolegend | 109808; RRID:AB_313445 |
| PerCP-CD45.2 (104) | Biolegend | 109826; RRID:AB_893349 |
| PE/Cy7-CD45.2 (104) | Biolegend | 109830; RRID:AB_1186098 |
| PB-CD45.2 (104) | Biolegend | 109820; RRID:AB_492872 |
| PB-CD62L (MEL-14) | Biolegend | 104424; RRID:AB_493380 |
| PE/Dazzle-CD64 (X54-5/7.1) | Biolegend | 139320; RRID:AB_2566559 |
| FITC-CD69 (H1.2F3) | Biolegend | 104506; RRID:AB_313109 |
| APC/Cy7-CD69 (H1.2F3) | Biolegend | 104526; RRID:AB_10679041 |
| FITC-CD8a (53-6.7) | Biolegend | 100706; RRID:AB_312745 |
| AF700-CD8a (53-6.7) | Biolegend | 100730; RRID:AB_493703 |
| PE/Cy7-F4/80 (BM8) | Biolegend | 123114; RRID:AB_893478 |
| PB-Granzyme B (GB11) | Biolegend | 515408; RRID:AB_2562196 |
| FITC-H-2kb (AF6-88.5) | BD Biosciences | 562002 |
| APC-H-2Ld MuLV gp70 Tetramer (N/A) | MBL International | TB-M521-2 |
| FITC-Isotype Rat IgG1, lambda (G0114F7) | Biolegend | 401913 |
| FITC-Isotype Rat IgM, kappa (R4-22) | BD Biosciences | 553942 |
| PE-Ki67 (11F6) | Biolegend | 151210; RRID:AB_2716008 |
| PerCP-Ly-6G (1A8) | Biolegend | 127654; RRID:AB_2616999 |
| FITC-MHCII (I-A/E) (M5/114.15.2) | Biolegend | 107606; RRID:AB_313321 |
| PB-MHCII (I-A/E) (M5/114.15.2) | Biolegend | 107620; RRID:AB_493527 |
| PE-Mouse IgG2a, kappa (MOPC-21) | Biolegend | 400113 |
| PE/Cy7-Mouse IgG2a, kappa (MOPC-21) | Biolegend | 400125 |
| PE-PD-1 (29F.1A12) | Biolegend | 135206; RRID:AB_1877231 |
| FITC-PD-1 (29F.1A12) | Biolegend | 135214; RRID:AB_10680238 |
| APC-PD-1 (RMP1-30) | Biolegend | 109112; RRID:AB_10612938 |
| APC-PD-1 (29F.1A12) | Biolegend | 135210; RRID:AB_2159183 |
| APC/Cy7-PD-1 (29F.1A12) | Biolegend | 135224; RRID:AB_2563523 |
| PE/Cy7-PD-L1 (10F.9G2) | Biolegend | 124314; RRID:AB_10643573 |
| PE-PD-L1 (10F.9G2) | Biolegend | 124308; RRID:AB_2073556 |
| BV421-PD-L1 (10F.9G2) | Biolegend | 124315; RRID:AB_10897097 |
| BV785-PD-L1 (10F.9G2) | Biolegend | 124331; RRID:AB_2629659 |
| PE/Dazzle-PD-L1 (10F.9G2) | Biolegend | 124324; RRID:AB_2565639 |
| APC-PD-L2 (TY25) | Biolegend | 107210; RRID:AB_2566345 |
| PE-PD-L2 (TY25) | Biolegend | 107206; RRID:AB_2162011 |
| PE-Perforin (S16009B) | Biolegend | 154406; RRID:AB_2721641 |

| | | |
|---|---|---|
| APC-pSTAT1 (S727) (A15158B) | Biolegend | 686408; RRID:AB_2650782 |
| APC-Rat IgG2a, kappa (RTK2758) | Biolegend | 400512 |
| PE/Cy7-Rat IgG2a, kappa (RTK2758) | Biolegend | 40522 |
| APC/Fire-Rat IgG2a, kappa (RTK2758) | Biolegend | 400567 |
| PE-Rat IgG2b, kappa (RTK4530) | Biolegend | 400636 |
| PerCP-Rat IgG2b, kappa (RTK4530) | Biolegend | 400630 |
| PE-SHP-2(pY542) (L99-921) | BD Pharmingen | 560389 |
| AF647-T-bet (4B10) | Biolegend | 644804; RRID:AB_1595466 |
| PE-TNFa (MP6-XT22) | Biolegend | 506306; RRID:AB_315427 |
| Mouse monoclonal anti-Stat1 (pY701) | BD Biosciences | 612133; RRID: AB_399504 |
| InVivoPlus Anti-mCD3e (145-2C11) | BioXcell | BP0001-1 |
| InVivoPlus mIgG (MOPC-21) | BioXcell | BP0083 |
| Mouse monoclonal anti-Stat1 | BD Biosciences | RRID: AB_397585 |

### Table S2. Reagents

| | | |
|---|---|---|
| **Recombinant Proteins and reagents** | | |
| mIL-6 | Biolegend | 575706 |
| mTNFα | BD Biosciences | 554589 |
| mGM-CSF | Biolegend | 576306 |
| LPS | | |
| mIFNα | R&D | 12100-1 |
| mIFNβ | Biolegend | 576306 |
| mIFNγ | Biolegend | 575306 |
| PMA | | |
| Ionomycin | | |
| Formalin, 10% (Phosphate Buffer). | ThermoFisher | SF100-4 |
| Zombie UV | Biolegend | 423108 |
| Mouse FcBlock | Biolegend | 101320 |
| mPD-L1-FC | Biolegend | 758208 |
| Fixation/Permeabilization Solution Kit with BD GolgiPlug | BD Biosciences | 555028 |
| HMW poly(I:C) | Invivogen | tlrl-pic |
| **Deposited Data** | | |
| BMS038: Tumor and Microenvironment Evolution during Immune Checkpoint Blockade Therapy with Nivolumab | Riaz et al., 2017 | https://github.com/riazn/bms038_analysis |
| Biomarkers of response and resistance to checkpoint blockade immunotherapy in metastatic melanoma | Gide TN, et al., 2019 | PRJEB23709 |
| Single-cell RNA-seq of melanoma ecosystems reveals sources of T cell exclusion linked to immunotherapy clinical outcomes | Jerby-Anon, et al. 2018 | Study: Melanoma immunotherapy resistance, Broad Single-Cell Portal |
| RNA seq analysis of SHP099 treated Leukemic Cells Expressing Genetic and Epigenetic Mutations | Pandey et al., 2019 | GSE134843 |
| Single cell transcriptomics of human and mouse lung cancers reveals conserved myeloid populations across individuals and species | Zilionis et al., 2019 | GSE127465 |
| **Oligonucleotides** | | |
| *Cd274*-sgRNA | Genscript | TCCAAAGGACTTGTACGTGG |
| Control-sgRNA | Genscript | CTCGTATCTTTTCCCACGGC |
| *B2m*-sgRNA | Genscript | AGTATACTCACGCCACCCAC |
| *Pdcd1*-sgRNA | Genscript | CAGCTTGTCCAACTGGTCGG |
| Mulv-gp70-FW | IDT | TGACCTTGTCCGAAGTGACC |
| Mulv-gp70-RV | IDT | TAGGACCCATCGCTTGTCTT |
| *Actb*-FW | IDT | ATTGTTACCAACTGGGACGACATG |
| *Actb*-RV | IDT | CTTCATGAGGTAGTCTGTCAGGTC |

**Table S3. Gene Signatures**

| SHP-2 Activation | $M_A$ | $M_B$ | $M_C$ | tPD-L1 Loss |
|---|---|---|---|---|
| NAT8L | MMP12 | SLC40A1 | CXCL10 | GPR141 |
| SERPINA3 | HAMP | EGR1 | CXCL11 | CCL2 |
| RPRM | GSDME | CD209 | GPR84 | MARCO |
| ABCA4 | TBC1D2 | IL2RA | ACOD1 | CCL8 |
| FCER1A | OSCAR | NR4A2 | GBP1 | DNASE1L3 |
| TLR7 | EIF4EBP1 | FOSB | G0S2 | JCHAIN |
| TUSC1 | PPARG | GPR34 | CXCL8 | TNFAIP2 |
| LDHB | PGD | NR4A1 | CCRL2 | EPSTI1 |
| LPCAT2 | UBASH3B | ARL4C | BCL2A1 | SOCS1 |
| GBP5 | NAGLU | RBP1 | NFKBIE | SLCO4A1 |
| DHRS1 | SMIM25 | RARRES1 | ISG15 | MSR1 |
| RNF128 | ACAA2 | EIF4A3 | CALHM6 | SLC28A2 |
| GAL | ARRDC4 | IQGAP2 | PLEK | GABRP |
| TMEM176B | MGLL | MERTK | TFEC | SCUBE3 |
| SDSL | PTPMT1 | EPHX1 | RNASE2 | CYP1A1 |
| RNASE6 | USF2 | ICAM4 | USP18 | C1QB |
| NLRP6 | ATP6V1C1 | MAF | RSAD2 | CCR9 |
| FCER1G | SLC16A6 | LILRB5 | METRNL | IRF7 |
| SLC6A12 | TIMM8B | GATM | SLC2A6 | TAP2 |
| SCIN | LAMTOR2 | IGF1 | MIR155HG | HLA-DQB1 |
| HOPX | TNS1 | MYO5A | UBE2L6 | HLA-DQB2 |
| MS4A2 | PNPO | IL13RA1 | IFIT1 | CD72 |
| SIRPB1 | MYO9B | PHACTR1 | CMPK2 | DCK |
| SIRPG | NOP10 | EPAS1 | OAS3 | SIGLEC1 |
| SIRPA | SPG21 | ADORA3 | TNFSF13B | MTHFD2 |
| CTSS | DENND4C | TMIGD3 | CCR5 | HMGA1 |
| TMEM53 | RAB10 | STOM | CD300E | PCDHB6 |
| SLC39A2 | NARS | HES1 | MYD88 | TAP1 |
| YPEL3 | ADCY3 | KCNJ5 | SERPINB2 | PSMB8 |
| GSTM1 | SSR3 | ITPR2 | OAS2 | PSMB9 |
| LCN2 | FDX1 | LINC01094 | B3GNT5 | MMP25 |
| TNFRSF18 | CD151 | CLIC2 | VAMP5 | CMTM7 |
| TNFSF12 | COPRS | SLC18B1 | PTGS2 | RNASE6 |
| HLA-DOA | PARP12 | CD72 | SERPINB9 | UBD |
| PDZK1IP1 | RPN1 | CD302 | APOL4 | SYCP2 |
| C12orf57 | SNX3 | CR1 | MS4A14 | PPA1 |
| TGFBR3 | ATP2C1 | CYBRD1 | VASH1 | IL18BP |
| CYP11A1 | ARHGDIA | CABLES1 | FCN1 | BATF2 |
| BPIFC | MTCH2 | MGAT4A | REL | AIF1 |
| CPQ | IRF5 | PXDC1 | RIPK2 | SLC39A2 |
| CSTA | PFKL | DRAM2 | SNAI1 | CALHM6 |
| CD27 | PSMA7 | TNFRSF21 | IL15RA | NRGN |
| SUOX | VTI1B | HERPUD1 | VDR | CCR5 |
| NDRG2 | GMFB | SNX9 | RBM47 | STAT1 |

| | | | | |
|---|---|---|---|---|
| TULP3 | RDX | C1orf54 | PARP14 | CYP2F1 |
| CMA1 | MDH1 | SERPINF1 | ARHGAP31 | PET100 |
| TMEM86A | HOMER3 | MAN1A1 | SMCO4 | BST2 |
| MGST2 | TOMM40 | GADD45G | GBP5 | USP18 |
| ASPRV1 | GTF3C6 | MTSS1 | TLR2 | USP41 |
| MAP1LC3A | TMED10 | EGR2 | ARHGEF2 | TMPRSS11E |
| CSRP3 | SLC25A39 | NAIP | IFIT3 | CDK1 |
| PCP4L1 | COMMD9 | SNX6 | SAMD9L | HTR2C |
| TP53INP2 | SLFN11 | MAN1C1 | FCGR3A | IRGM |
| TMTC2 | ME2 | MGST2 | TRAF1 | IFIT3 |
| IL1RN | PPIB | IFNGR1 | IFI44 | TNS4 |
| MS4A7 | VAMP3 | NEU1 | SPHK1 | MCM5 |
| UBE2L6 | SLC39A1 | IER2 | ACSL1 | SLFN13 |
| TYROBP | PDCD6IP | GAS6 | GLRX | UHRF1 |
| GP1BB | MAN2B2 | FOS | IFNGR2 | GBP3 |
| CYP4V2 | NDUFB9 | RANBP2 | RAPGEF2 | OAS2 |
| TPSG1 | RHEB | SLC46A3 | PSTPIP2 | SAA2 |
| PDK2 | APH1A | TPCN1 | PITPNA | SAA1 |
| CX3CR1 | YWHAG | SCAMP4 | GNA13 | APOL6 |
| LIF | GGA2 | LINC00847 | SNX8 | MS4A6A |
| CSTB | EHD4 | ST3GAL1 | TET2 | MS4A6E |
| C15orf48 | ADAM17 | GBGT1 | XAF1 | CXCL9 |
| MGST1 | ALDH3B1 | RALGDS | PTPRE | GBP6 |
| TMEM216 | SEC61A1 | CITED2 | ATP2B1 | MS4A7 |
| RAB38 | VPS35 | AC009404.1 | MTHFD2 | DRAXIN |
| GSTK1 | HADHA | WDR45B | PGAM1 | TOP2A |
| RRAS | UQCR11 | ITSN1 | KDM6B | KLK1 |
| TCN2 | GLUD1 | TNFSF15 | COL8A2 | CDCA8 |
| VAT1L | NRAS | LATS2 | PLEKHO1 | SLAMF8 |
| MAOB | HDLBP | RBMS1 | IFITM3 | ISG15 |
| CLNK | ELOB | PRMT9 | RNASEK | GBP2 |
| BST1 | USMG5 | JUNB | ADPGK | MX1 |
| HS3ST1 | AUP1 | ATF6 | STAT2 | LCN2 |
| SLC30A2 | REEP5 | ANKH | CDCP1 | CTLA4 |
| FCGRT | ATP5F1 | CCNQ | TNIP3 | CCNA2 |
| PSAP | PPDPF | SLC38A7 | RAPGEF1 | DHX58 |
| RAB19 | GALNT1 | SUMF2 | IFIT2 | MNDA |
| IFI27L2 | VMA21 | PLOD1 | IFI35 | RSAD2 |
| DENND2A | ATP5I | TSC22D1 | ACKR3 | SMPDL3B |
| ICAM4 | KDELR2 | METTL7A | NFKBID | CXCL13 |
| PRDX5 | ST13 | NFIL3 | GBP4 | FCGR1A |
| LST1 | ERO1A | HSPH1 | MAP2K1 | OAS1 |
| P2RX4 | CD52 | GOLIM4 | BASP1 | GP9 |
| HEBP1 | PPIA | WIPI1 | PSMA4 | OAS3 |
| SLC9A9 | | ANKRD9 | CD44 | S100A4 |
| PGLYRP1 | | AL033543.1 | ZFYVE16 | IGKC |
| FFAR2 | | SNX29 | FKBP1A | IFIT1B |

| | | | | |
|---|---|---|---|---|
| LYPD6 | | RCAN1 | FLT1 | SCGB1A1 |
| CIB2 | | HSD17B12 | YWHAE | SCGB3A2 |
| TMEM215 | | ID2 | HCST | SCIMP |
| F10 | | TRPS1 | CTNNA1 | ZBP1 |
| CD52 | | UGCG | FUOM | FCGR3A |
| BCAS1 | | ZBED1 | LPCAT2 | FCGR3B |
| ORM2 | | CHID1 | RABGEF1 | NLRC5 |
| ORM1 | | SESTD1 | FAM102B | P2RY12 |
| PTGES | | IER5 | CYCS | IDO1 |
| GNG11 | | TM7SF3 | FAM49A | S100A8 |
| TMEM158 | | XIST | TMEM109 | CXCL11 |
| HEXB | | HNRNPU | CEBPB | XAF1 |
| SMIM4 | | SPATS2L | PSMB2 | RTP4 |
| FUOM | | CD93 | MAP4 | OASL |
| GNGT2 | | C3 | ARPC4 | CXCL10 |
| HCK | | | WSB2 | POLE |
| CLU | | | ETV3 | DCTPP1 |
| NAAA | | | ACSL4 | P2RY13 |
| MCEMP1 | | | WDFY1 | IFI44 |
| CAMP | | | MTF1 | PHF11 |
| KLRB1 | | | DENND5A | CD33 |
| VCAM1 | | | ATP5J2 | SIGLEC6 |
| NUPR1 | | | ECHS1 | FCGR1B |
| MSRB1 | | | TWF2 | CD274 |
| CD302 | | | CFLAR | FCMR |
| PODXL | | | EIF2AK2 | HSPBAP1 |
| DAPK2 | | | VMO1 | LAIR1 |
| LRG1 | | | SLC2A14 | KIF2C |
| MYL10 | | | DOCK10 | LGALS3BP |
| GPX3 | | | GCH1 | S100A9 |
| S100A8 | | | ERP44 | PPBP |
| CD63 | | | CCT5 | IGHM |
| CPA3 | | | RCC2 | |
| RGS2 | | | TPM3 | |
| ANKRD22 | | | KLF10 | |
| NREP | | | ARL5B | |
| SMIM19 | | | PTPA | |
| S100A6 | | | SSB | |
| RMRP | | | PTBP1 | |
| MOGAT2 | | | IRF7 | |
| DNAJA4 | | | APP | |
| TMEM9 | | | DDX3Y | |
| SLC18A2 | | | EAF1 | |
| HMGN3 | | | PDE4A | |
| FPR3 | | | CYC1 | |
| FPR2 | | | TLNRD1 | |
| FKBP1B | | | EHD1 | |

| | | | |
|---|---|---|---|
| TARM1 | | | HMGA1 |
| MAP6 | | | HMGN2 |
| TNFSF13B | | | RRBP1 |
| NKG7 | | | NCOR2 |
| CHIA | | | PPA1 |
| HPR | | | MT1G |
| HP | | | TESK1 |
| ISOC2 | | | C5orf15 |
| SMIM24 | | | SLC2A3 |
| | | | SLC39A8 |
| | | | PLXNB2 |
| | | | ARFGAP3 |
| | | | OGFOD1 |
| | | | HPSE |
| | | | HSP90AB1 |
| | | | BTG3 |
| | | | LYPD3 |
